# Holographic stimulation of opposing amygdala ensembles bidirectionally modulates valence-specific behavior

**DOI:** 10.1101/2022.07.11.499499

**Authors:** Sean C Piantadosi, Zhe Charles Zhou, Carina Pizzano, Christian E Pedersen, Tammy K Nguyen, Sarah Thai, Garret D Stuber, Michael R Bruchas

## Abstract

The basolateral amygdala (BLA) is an evolutionarily conserved brain region, well known for valence processing. Despite this central role, the relationship between activity of BLA neuronal ensembles in response to appetitive and aversive stimuli and the subsequent expression of valence-specific behavior has remained elusive. Here we leverage 2-photon calcium imaging combined with single cell holographic photostimulation through an endoscopic lens implanted in the deep brain to demonstrate a direct causal role for discrete ensembles of BLA neurons in the control of oppositely valenced behavior. We report that targeted photostimulation of individual groups of appetitive or aversive BLA neurons shifts behavioral responses toward those behaviors which recruited a specific consumption or avoidance ensemble. Here we identify that neuronal encoding of valence in the BLA is graded and relies on the relative proportion of individual BLA neurons recruited in a stable appetitive or aversive ensemble.

## Introduction

To ensure survival, organisms must rapidly and accurately assess the safety of their environment and various stimuli. Some stimuli are innately aversive, such as bitter tastes indicating an item is unsafe to consume, while others are innately appetitive, such as sugar rich sweet tasting foods. The perceived value of these stimuli, also known as valence, dictates subsequent approach or avoidance behavior. The basolateral amygdala (BLA) has been identified as a critical node where specific information about stimulus valence is encoded (Beyeler et al., 2018; Gore et al., 2015; Grewe et al., 2017; Kyriazi et al., 2018; Morrison and Salzman, 2011; Paton et al., 2006; Zhang and Li, 2018). However, prior experimental and theoretical exploration of valence coding within the BLA has relied on indirect correlations of neuronal activity with behavior (Beyeler et al., 2018; Borst and Theunissen, 1999; Grewe et al., 2017; Gründemann et al., 2019; Kumar et al., 2010).

Within the BLA, spatially intermixed groups of genetically similar excitatory neurons (However see (Kim et al., 2016)) are known to respond to either appetitive or aversive unconditioned stimuli (US) with innate valences (Beyeler et al., 2018; Gore et al., 2015; Zhang and Li, 2018), and this response selectivity can be imbued on previously neutral stimuli that predict the US (Grewe et al., 2017). However, conventional optogenetic approaches to causally probe neural activity lack spatial specificity to target specific intermixed neural ensembles in freely behaving animals; thus the direct causal role of opposing BLA neuronal ensembles in driving valence specific behavior has yet to be established. Moreover, it is unknown whether the activity of these opposing ensembles contribute equally to valence-specific behavior generation (Tye, 2018).

Identifying the key features of deep brain neural responses which encode valence information to initiate consumption and avoidance remains a fundamental question. Existing approaches have been used to manipulate distinct groups of neurons in the BLA have relied on immediate early gene expression (Gore et al., 2015), which lacks the sensitivity to detect decreases in neural activity, provides slow and imprecise labeling, and labels neurons that respond to both appetitive and aversive stimuli, which constitute a large population of BLA neurons (Shabel and Janak, 2009). To ascertain whether the expression of valence-specific behaviors results from the activity of discrete BLA ensembles, a method for manipulating the activity of these ensembles in a temporally and spatially precise manner is necessary.

Recent advances in multiphoton imaging, opsin development, and targeted photo-stimulation technology has afforded investigators the ability to conduct simultaneous recording and manipulation of neural activity at the single cell level (Adesnik and Abdeladim, 2021; Carrillo-Reid et al., 2017, 2019; Daie et al., 2021; Gill et al., 2020; Jennings et al., 2019; Marshel et al., 2019; Robinson et al., 2020; Russell et al., 2022; Sridharan et al., 2022; Yang et al., 2018). This convergence of technology has led to the ability to directly define causality of discrete neuronal ensembles in mediating a multitude of behaviors (Carrillo-Reid et al., 2019; Daie et al., 2021; Gill et al., 2020; Robinson et al., 2020; Yang et al., 2018). However, recent investigations using this method have focused on superficial cortical regions (Carrillo-Reid et al., 2019; Gill et al., 2020; Jennings et al., 2019; Marshel et al., 2019) as well as the hippocampal formation(Robinson et al., 2020). Ensemble coding in these brain regions, while stable, is often distributed, highly stochastic, and subject to representational drift (Pérez-Ortega et al., 2021; Rule et al., 2019; Ziv et al., 2013). To date, no experiments have harnessed this technology for interrogation of deep brain (Adesnik and Abdeladim, 2021), evolutionarily conserved limbic structures, like the BLA, well known to encode stimulus valence. Here we determined whether valence is stably encoded at the level of distinct separable groups of single neurons within the BLA, and whether valence-specific changes in neuronal activity are sufficient to bidirectionally control consumption or avoidance behavior.

## Results

### Optically separable populations of BLA neurons respond to appetitive and aversive stimuli

To determine how individual principal neurons within the BLA encode opposing valences, we developed an optical approach to simultaneously monitor as well as modulate the activity of individual BLA neurons through a gradient refractive index lens (GRIN). To achieve this, a combination (3:1 ratio) of two adeno-associated viruses encoding either the calcium indicator GCaMP6f (AAV5-CaMKIIa-GCaMP6f) and the red-shifted excitatory opsin ChRmine (AAV8-CaMKIIa-ChRmine-mScarlet-Kv2.1-WPRE) were co-injected unilaterally into the BLA of wildtype (C57BL/6) mice (**Figure 1A**). A GRIN lens was implanted directly above our viral injection target. Colocalization of GCaMP6f and ChRmine was observed within the BLA underneath the GRIN lens (**Figure 1B; Figure S1A**). Post-hoc confirmation of GRIN lens placement within the BLA was conducted for each mouse (GRIN lenses spanned -1.46 to -1.94mm from Bregma; **Figure S1B**).

**Figure 1.**
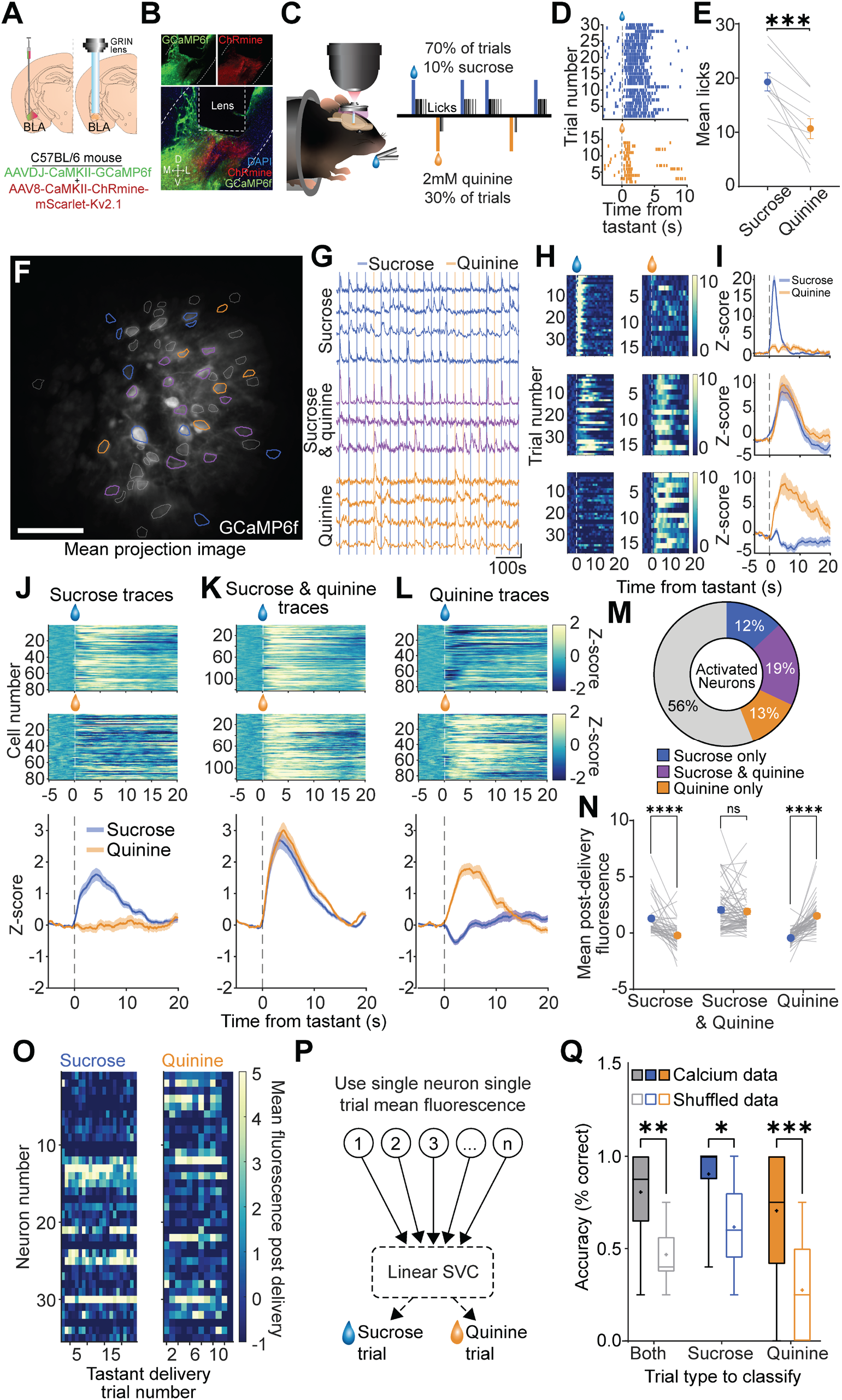
Optically separable populations of BLA neurons respond to appetitive and aversive stimuli. (**A**) Surgical schematic demonstrating co-injection of viral vector to express GCaMP6f and ChRmine in the same BLA neurons and subsequent GRIN lens implantation. (**B**) Histological image of co-expression of GCaMP6f (green) and ChRmine (red) in the BLA just below GRIN lens. (**C**) Schematic depiction of head-fixed mouse under the 2-photon microscope and breakdown of trial distribution. On 70% of trials, 10% sucrose (blue) was delivered, while on 30% of trials 2mM quinine (orange) was delivered (20-25s ITI). Example licking rasters aligned to tastant delivery from a single sucrose and quinine delivery session separated by trial type (top – sucrose, bottom – quinine). Lines indicate individual licks. (**E**) Mean number of licks on sucrose versus quinine delivery trials across mice (*t*(9)=6.14,*p*=0.0002). (**F**) Mean projection image from representative FOV of BLA neurons during sucrose and quinine session. Neurons are colored according to responsivity to tastants. Grey=unaffected, Blue=sucrose activated, Orange=quinine activated, Purple=sucrose and quinine activated. (**G**) Representative continuous traces of neurons colored according to **F** overlayed on top of individual tastant delivery trials (Sucrose=blue, Quinine=orange). (**H**) Representative heatmaps of tastant-delivery aligned calcium fluorescence from individual neurons during sucrose (left) or quinine (right) delivery trials for a sucrose activated (top), sucrose and quinine activated (middle), or quinine activated (bottom) neuron. (**I**) Calcium fluorescence averaged across trials for fluorescence data in (**H**). (**J**) Trial-averaged fluorescence of sucrose activated neurons in response to sucrose (top) or quinine (middle) delivery. Mean response across all neurons on sucrose (blue) or quinine (orange) delivery trials (bottom). (**K**) Trial-averaged fluorescence of sucrose and quinine activated neurons in response to sucrose (top) or quinine (middle) delivery. Mean response across all neurons on sucrose (blue) or quinine (orange) delivery trials (bottom). (**L**) Trial-averaged fluorescence of quinine activated neurons in response to sucrose (top) or quinine (middle) delivery. Mean response across all neurons on sucrose (blue) or quinine (orange) delivery trials (bottom). (**M**) Proportions of unaffected (grey), sucrose activated (blue), sucrose and quinine activated (purple), or quinine activated (orange) neurons across entire population (*n*=651 neurons). (**N**) Mean fluorescence during post-tastant delivery period for sucrose activated, sucrose and quinine activated, and quinine activated neurons (Main effect of activated cell type [*F*(2,164)=21.76,*p*<0.0001], interaction between activated cell type and tastant delivery [*F*(2,164)=34.97,*p*<0.0001]. *Post-hoc* comparison of tastant delivery on sucrose activated neurons (*t*(164)=4.79,*p*<0.0001) and quinine activated neurons (*t*(164)=6.92,*p*<0.0001), no effect of tastant delivery type on sucrose and quinine activated neuron activity *p* > 0.05). (**O**) Population activity vectors from one mouse occurring during tastant consumption for sucrose (left; blue) and quinine (right; orange) trials. Each row represents the mean fluorescence for a single neuron on a single trial. (**P**) Schematic depicting binary linear support vector classifier (SVC) trained on single neuron mean fluorescence (**Q**) Accuracy of linear SVC for predicting either sucrose or quinine trials (both; black), only sucrose trials (blue), or only quinine trials (orange) across mice (*n*=13). Filled box-and-whisker plots indicate classifier trained on true fluorescence data. Unfilled box-and-whisker plot represents classifier trained on fluorescence traces that have been randomly shuffled in time. Whiskers represent minimum and maximum accuracy values for each individual mouse. + symbol denotes mean accuracy across mice. Both tastant (*t*(36)=3.35, *p*=0.006), sucrose only (*t*(36)=2.84, *p*=0.02), quinine only (*t*(36)=4.3, *p*=0.0004). \*\*\*\**p*<0.0001,\*\*\**p*<0.001.

We first established mice in a behavioral paradigm where they receive random presentations (ITI = 20-25s, total session duration = 20 min) of either un-cued 10% sucrose (sucrose) or 2mM quinine (quinine), highly appetitive and aversive tastants, respectively. To ensure consistent consumption of both tastants, trial presentation was structured such that on 70% of trials sucrose was delivered and on 30% of trials quinine was delivered (**Figure 1C**) (K Namboodiri et al., 2021). The un-cued nature of delivery combined with limited water restriction ensured that mice continued to stably sample each tastant by licking throughout the duration of an entire session (**Figure 1D**). During a session in which 100% of trials resulted in sucrose delivery, water restricted mice continued to lick consistently with no signs of satiety (**Figure S2A-D**). By contrast, when 100% of trials resulted in quinine delivery, water restricted mice ceased licking by the end of the session (**Figure S2C-D**) demonstrating its aversive quality. Mice licked significantly more on sucrose trials relative to a quinine trials (**Figure 1E**) Compared to trials in which drinking water was delivered (**Figure S2E**), licking frequency was significantly elevated when sucrose was delivered and significantly reduced on quinine trials (**Figure S2F-H**). During these behavioral sessions the activity of individual excitatory neurons expressing GCaMP6f within the BLA were recorded (total = 651 neurons, mean neurons per mouse = 46.5 *n*=14 mice) (**Figure 1F**). We compared individual trials of neural activity to shuffled versions of the same activity traces, maintaining identical variance, to classify neurons as either only sucrose activated, both sucrose and quinine activated, or only quinine activated (**Figure 1G**). Individual neural responses for significantly responsive neurons were robust and consistent to their preferred tastant across trials, and sucrose- or quinine-only activated neurons displayed negligible responses to the unpreferred tastant (**Figure 1H-I**). This same activity pattern was observed in the trial-averaged data across mice, with sucrose-activated neurons displaying robust increases in activity in response to sucrose, but not quinine (**Figure 1J**), sucrose and quinine-activated neurons exhibiting similar increases in activity in response to both the appetitive and aversive tastant (**Figure 1K**), and quinine neurons showing a robust increase in activity specifically to quinine (**Figure 1L**). Large groups of neurons that were inhibited in response to sucrose-only, sucrose and quinine, or quinine-only were also identified (**Figure S3A-C**). Interestingly, many neurons displayed seemingly opposing activity changes, with increases or decreases in activity to sucrose associated with an opposing change in response to quinine (**Figure S3E-F**). Despite differences in licking behavior between the appetitive and aversive tastant (**Figure 1D-E, Figure S2**), no differences in the mean fluorescence traces of sucrose relative to quinine activated neurons were observed (**Figure S3G**), suggesting that these neural responses are linked to valence rather than to differences in consumption or different levels of GCaMP6f expression in the neurons. Across the entire population of BLA neurons, 19% of neurons (122/650) were identified as sucrose and quinine activated neurons, while 12% (81/650) and 13% (85/650) of neurons were sucrose only and quinine only activated, respectively (**Figure 1M**). The mean calcium fluorescence for each neuron following tastant delivery (0 to 10s) showed strong and consistent increases in fluorescence for sucrose-activated neurons and quinine-activated neurons to their preferred trial conditions respectively, while no difference in fluorescence was detected for neurons classified as both sucrose and quinine activated (**Figure 1N**). We believe this response reflects stimulus valence rather than a difference in licking structure, as the mean fluorescence for each sucrose- or quinine-activated neuron was significantly different when analysis was restricted to the immediate post-delivery period where licking behavior was not different between trial types (**Figure S3H-J**). We note that we did not identify any differences in the anatomical location (medial/lateral or anterior/posterior) of sucrose activated neurons and quinine activated neurons were detected across all recorded neurons (**Figure S3K-L**). Single neuron responses to sucrose or quinine were also stable across multiple sessions (**Figure S4A-F**) both in terms of amplitude of the response (**Figure S4G**) and the temporal dynamics of the fluorescence signal (**Figure S4H-I**). Finally, using the mean fluorescence response for each neuron on each tastant delivery trial (**Figure 1O**) we trained a binary linear support vector classifier (SVC), ideally suited for two-class problems, to predict tastant trial type (**Figure 1P**). Mean BLA population activity in the post-tastant epoch reliably predicted whether sucrose or quinine were delivered compared to a model trained on shuffled calcium fluorescence data, containing equal mean fluorescence and variance as the unshuffled calcium fluorescence. Classifier accuracy was also significantly better than shuffled control data for both individual tastant (sucrose or quinine) delivery trials (**Figure 1Q**). These results suggest that the activity of groups of individual BLA neurons stably encode the valence of a stimulus.

### Holographic photostimulation via a GRIN lens is specific and robust at activating amygdala ensembles

Having established that intermixed but spatially separable populations of BLA neurons stably encode stimulus valence we tested whether these correlative changes in activity were sufficient to drive corresponding appetitive and aversive behavioral responding. We used a spatial light modulator (SLM) (Adesnik and Abdeladim, 2021; Carrillo-Reid et al., 2017) to photostimulate predefined neurons in the field of view (**Figure 2A**) expressing the red-shifted excitatory opsin ChRmine (Marshel et al., 2019) (**Figure 2B**). We first calibrated and determined the specificity and physiological resolution tolerance of single-neuron stimulation in the X/Y plane. To achieve this, we targeted a single neuron in our field of view expressing both GCaMP6f and ChRmine for spiral stimulation (10hz, 2s, 8mW power per ROI). We then proceeded to move the spiral stimulation target away from the cell body at a distance of 10um each trial (roughly 1 cell-body width, 5 trials per ROI, randomized) for 5 sequential stimulation targets up to 40um away from the targeted neuron (**Figure 2C**). We found that photoactivation of originally targeted neuron fell off dramatically as the stimulation ROI was moved laterally (**Figure 2D**), such that once the spiral stimulation target was > 1 cell body width (∼20µm) away there was no change in fluorescence of the original neuron (**Figure 2E**), indicating a very narrow window of control over the x/y cartesian plane at the single neuron level. We then conducted a subsequent experiment to determine the resolution specificity of stimulation in the Z-axis by first identifying neurons for stimulation in a recording plane and then, for different trials, stimulating at various planes above and below the recording plane. To record the effect of out of plane stimulation on the fluorescent activity of the target neuron in the recording plane, we coupled an electrically tunable lens (ETL) to the microscope imaging pathway which was used to near-simultaneously (< 5ms) compensate for the change in microscope Z-position to image the recording plane while maintaining the photostimulation z-depth location (**Figure 2F**). We next measured and corrected for the amount magnification caused by the GRIN lens at varying depths (**Figure 2G; Figure S5A-C**). Using this method, we established a clear distance-response relationship, with stimulation targeted to the recording plane (±0µm) eliciting the greatest change in fluorescence of the target neuron (**Figure 2H**). At the extremes of the microscope Z-position movements (-26 and +33µm in converted distance) limited activation of the target neuron (*n*=6 neurons) was seen relative to stimulation within the recording plane across 5 individual trials (**Figure 2I**). Across all distances, we found a similar relationship between distance and stimulation evoked response to what we observed in **Figure 3D**, with the effects of out of plane photostimulation negligible at distances of 20µm above or below the imaging plane (**Figure 2J**). Finally, we find that deep brain SLM stimulation through a GRIN lens scales with the number of neurons targeted, allowing for robust activation of multiple neurons simultaneously with no change in the timing or amplitude of the photostimulated response as a function of neuron number (**Figure 2K**). Further, repeated stimulation of the same neurons produced consistently similar increases in fluorescence and did not decrease in magnitude over the course of a session (**Figure 2L**). Together, these results establish specificity of photostimulation through a GRIN lens within the BLA and a working paradigm for targeted, repeatable control of individual neuron activity using SLM deep in the brain of an awake, behaving mouse. These findings demonstrate holographic SLM stimulation through a deep brain GRIN lens, like the BLA as capable of robust, precise, and consistent activation of a priori optically identified neurons.

**Figure 2.**
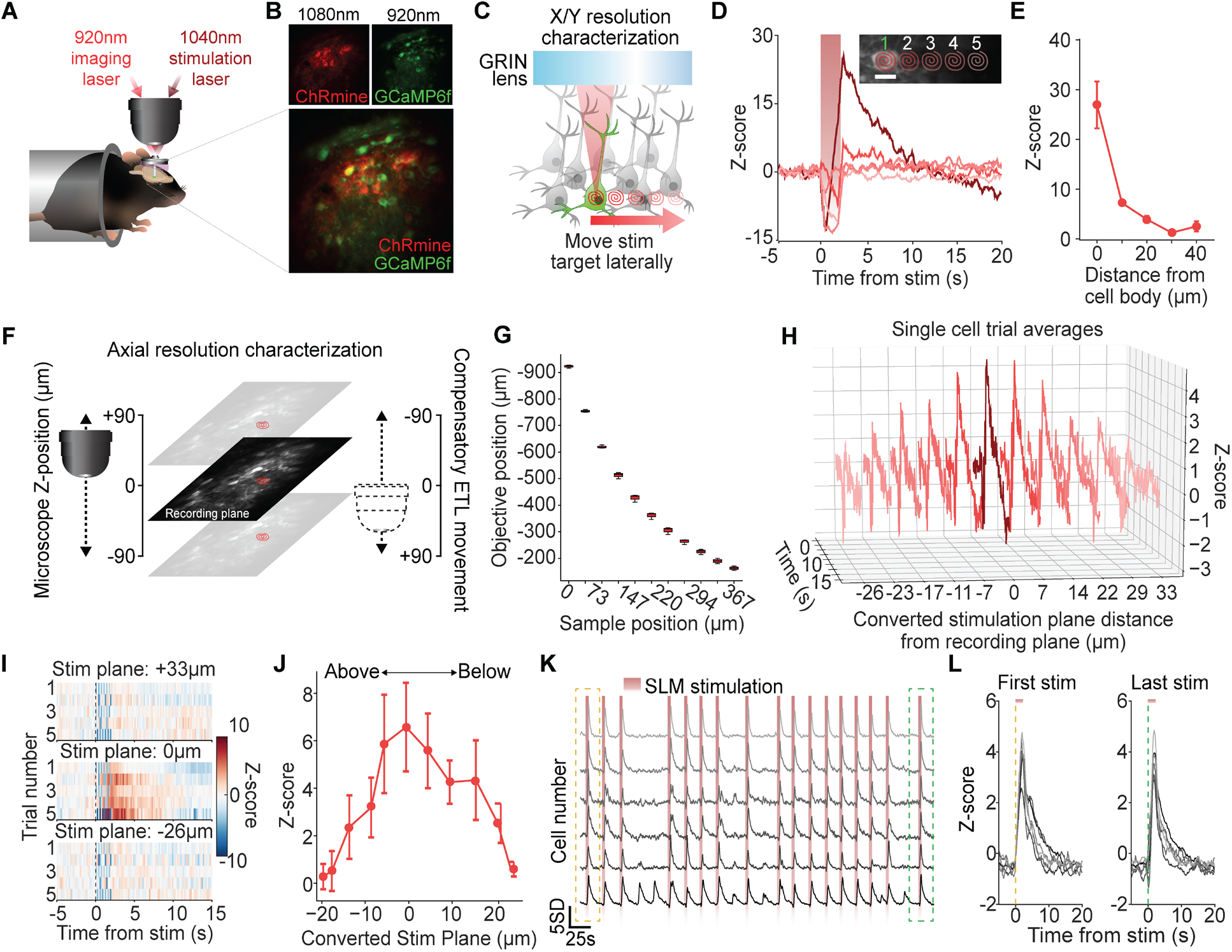
Deep brain holographic stimulation of individual BLA neurons is specific and robust. (**A**) Schematic of head-fixed 2-photon calcium imaging via 920nm laser path combined with a second 1040nm laser path for stimulation. (**B**) *In vivo* imaging of GCaMP6f (920nm) or the red-shifted opsin ChRmine (1080nm) expressed in BLA neurons. (**C**) Schematic depicting experimental setup for testing X/Y specificity of single ROI activation using a spatial light modulator (SLM). Red spirals indicate SLM targets, with each subsequent stimulation moving a fixed distance (+10, +20, +30, +40µm) laterally from the target cell. (**D**) Mean fluorescence across 5 stimulation trials (10hz, 2s, 6mW power) at each distance, darker colors indicate SLM spiral target being closer to the target cell. (Inset) Mean projection image of neuron recorded from (green) and spiral stimulation targets moving laterally. Order of spiral stimulation target (1-5) was randomized. (**E**) Mean fluorescence of single BLA neuron across 5 stimulation trials at each SLM ROI plotted as a function of distance from the target neuron. (**F**) Schematized representation of strategy to assess axial specificity of single-cell SLM stimulation. To stimulate (10hz, 2s, 6mW power) both above and below our target cell, the microscope Z-position was moved at fixed distances (-90 to +90µm in microscope space). At the same time, a separate imaging path with an electrotunable lens (ETL) was used to counteract this microscope movement to ensure imaging of the plane where our target cell was located. (**G**) Sample position and corresponding objective position for determining the degree of axial magnification caused by the GRIN lens. (**H**) Trial averaged fluorescence of a single BLA neuron plotted as a function of both time (left Y-axis) and stimulation distance in microscope units (X-axis). (**I**) Heatmaps showing stimulation aligned fluorescence traces (dotted line indicates onset of SLM stimulation) for 3 distances across 5 repeated stimulation trials (ITI=15s). (**J**) SLM stimulation evoked fluorescence averaged across all neurons (*n*=6) plotted as a function of distance corrected for GRIN lens magnification (brain distance in µm). Negative numbers indicate SLM stimulation ROIs above the target neuron, while positive numbers indicate SLM stimulation ROIs below. (**K**) Simultaneous activation of 6 BLA neurons through a GRIN lens using SLM stimulation (10hz, 2s, 6mW power per ROI). Yellow box indicates first stimulation of the session, green box indicates final stimulation of the session. (**L**) Aligned SLM stimulation evoked fluorescence from the first stimulation (left; yellow line) and the final (right; green line) stimulation. Traces are shaded according to their color in **K**.

**Figure 3.**
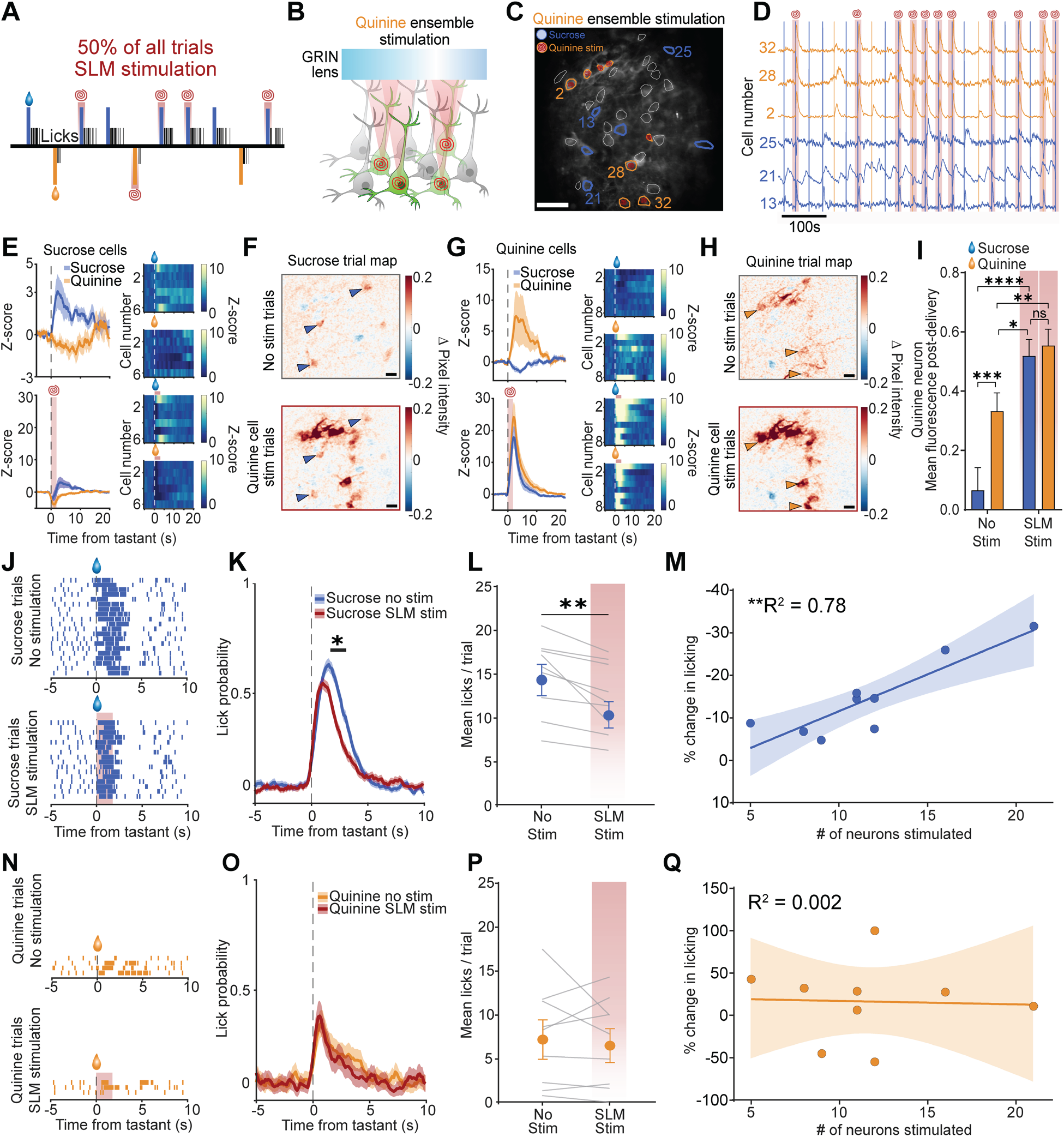
Selective simulation of aversive ensemble reduces licking of appetitive tastant. (**A**) Schematic of trial structure for SLM stimulation sessions. 70% of trials resulted in 10% sucrose (blue) delivery, while 30% of trials resulted in 2mM quinine (orange) delivery. SLM stimulation (10hz, 2s, 6-10mW power per ROI) was randomly delivered simultaneously with tastant delivery on 50% of all trials. (**B**) SLM stimulation ROIs were targeted to quinine-activated (quinine-ensemble) neurons. (**C**) Mean projection image of FOV from a single mouse. BLA neuron contours are colored according to their activity during the sucrose and quinine delivery session. Blue contours indicate neurons that were activated in response to sucrose only, while orange contours indicate neurons activated in response to quinine only. SLM spiral stimulation targets indicate the 8 quinine activated neurons (quinine ensemble) targeted for stimulation. (**D**) Example traces from 3 sucrose (blue) and quinine (orange) activated neurons labeled in **C**. Traces are overlayed on top of tastant delivery trials (sucrose=blue, quinine=orange) as well as trials in which SLM stimulation of the quinine ensemble occurred (red shading). Mean fluorescence activity of sucrose activated neurons in response to sucrose (blue) and quinine (orange) delivery on trials without stimulation (top) and trials with SLM stimulation (bottom). Dotted line indicates tastant delivery. Red shading indicates onset of quinine ensemble stimulation via SLM. (**F**) Spatial map of mean pixel intensity during the post-sucrose delivery period subtracted from mean pixel intensity immediately preceding sucrose delivery (-5s to 0s) during trials without quinine ensemble stimulation (top) and SLM stimulation of the quinine ensemble (bottom). Blue triangles indicate location of sucrose activated neurons identified in **C** and **D**. Red shading indicates a greater difference in activity during the post-sucrose delivery period relative to baseline. Blue shading indicates reduced activity. (**G**) Mean fluorescence activity of quinine activated neurons in response to sucrose (blue) and quinine (orange) delivery on trials without stimulation (top) and trials with SLM stimulation (bottom). Dotted line indicates tastant delivery. Red shading indicates onset of quinine ensemble stimulation via SLM. (**H**) Spatial map of mean pixel intensity during the post-quinine delivery period subtracted from mean pixel intensity immediately preceding quinine delivery (-5s to 0s) during trials without quinine ensemble stimulation (top) and SLM stimulation of the quinine ensemble (bottom). Orange triangles indicate location of quinine activated neurons identified in **C** and **D**. Red shading indicates a greater difference in activity during the post-quinine delivery period relative to baseline. Blue shading indicates reduced activity. (**I**) Mean fluorescence in the post-delivery period for quinine neurons in response to sucrose and quinine delivery on no stimulation versus SLM stimulation (two-way repeated measures ANOVA main effect of stimulation *p*<0.0001, main effect of tastant *p*<0.01, stimulation X tastant interaction *p*<0.01. Sidak’s multiple comparison correction displayed) (**J**) Lick rasters from a single mouse organized by trial type and aligned to sucrose delivery on trials without quinine ensemble SLM stimulation (top) and on trials with quinine ensemble stimulation (bottom; red shading). (**K**) Licking probability across all mice on sucrose delivery trials without stimulation (blue) and on trials with simultaneous quinine ensemble SLM stimulation (red) (one-way ANOVA, *p*<0.001). (**L**) The mean number of licks per trial per mouse on sucrose delivery trials without quinine ensemble stimulation and on trials with quinine ensemble SLM stimulation (red shading) (*t*(8)=3.73, *p*=0.006). (**M**) Correlation between the number of quinine ensemble neurons stimulated via SLM and the percent change in licking on sucrose trials (No stimulation vs quinine ensemble stimulation trials) (*R*^2^=0.78, linear regression *F*(1,7)=25.0 *p*=0.002). (**N**) Lick rasters from a single mouse organized by trial type and aligned to quinine delivery on trials without quinine ensemble SLM stimulation (top) and on trials with quinine ensemble stimulation (bottom; red shading). (**O**) Licking probability across all mice on quinine delivery trials without stimulation (blue) and on trials with simultaneous quinine ensemble SLM stimulation (red). (**P**) The mean number of licks per trial per mouse on quinine delivery trials without quinine ensemble stimulation and on trials with quinine ensemble SLM stimulation (red shading) (*t*(8)=0.35, *p*=0.73). (**Q**) Correlation between the number of quinine ensemble neurons stimulated via SLM and the percent change in licking on quinine trials (No stimulation vs quinine ensemble stimulation trials) (*R*^2^=0.002, linear regression *F*(1,7)=0.01 *p*=0.92). \*\**p*<0.01.

### Targeted photostimulation of aversive ensemble reduces consummatory behavior of appetitive stimulus

We next tested the central hypothesis for a direct causal link between these separable neural responses and behavior. We used a behavioral paradigm in which spiral SLM stimulation (2s, 10hz activation, power per ROI 5-10mW) occurred randomly on 50% of all tastant (35% sucrose trials and 15% quinine trials) delivery trials (**Figure 3A**). We first tested whether selective activation of the aversive, quinine-responsive ensemble altered licking behavior (**Figure 3B**). On average we targeted 10.7 quinine neurons per mouse (range 4 to 21/mouse, total=87) for SLM stimulation. We evaluated whether SLM stimulation of the quinine ensemble produced neuronal activity changes selectively in quinine activated neurons and not in sucrose activated neurons identified during baseline sessions (**Figure 4C-D**). On trials without holographic stimulation of the quinine ensemble, sucrose neurons maintained their preference for sucrose over quinine (**Figure 3E-F**; top, *n* = 6 neurons from representative mouse), consistent with their activity during baseline sessions (**Figure 1**). Critically, the activity of sucrose neurons was not affected by SLM stimulation of the quinine ensemble, with these neurons exhibiting a similar activity increase selectively to sucrose even during quinine ensemble stimulation trials (**Figure 3E-F**; bottom). By contrast, quinine-selective neurons displayed a typical preference for quinine delivery over sucrose on trials without SLM stimulation of the quinine ensemble (**Figure 3G-H**; top, *n* = 8 neurons from representative mouse). However, quinine ensemble neuron activity was not different on trials in which either sucrose or quinine was delivered concurrently with quinine ensemble activation (**Figure 3G-H**; bottom). Across all quinine neurons from all mice, the mean fluorescence in response to quinine delivery on trials without targeted photostimulation of the quinine ensemble was significantly elevated relative to sucrose delivery trials (**Figure 3I**). When SLM photostimulation of the aversive ensemble was paired with sucrose delivery, a significant increase in fluorescence was detected relative to the no stimulation sucrose delivery trials. We also observed a significant but modest increase in fluorescence of the quinine neurons when photostimulation was paired with quinine delivery, likely due to the difficulty of titrating laser power on an individual neuron basis. Importantly, no significant difference in mean fluorescence was observed between trials in which photostimulation was paired with sucrose or quinine, establishing specific “play back” of this aversive ensemble activity pattern on sucrose delivery trials in which there would typically have been no activity in quinine neurons. No change in fluorescence was detected when quinine activated neurons were targeted for SLM stimulation in mice not expressing ChRmine (**Figure S6A-D**). These data indicate that SLM activation of the aversive quinine ensemble was specific in both the spatial and temporal dimensions during our sessions.

**Figure 4.**
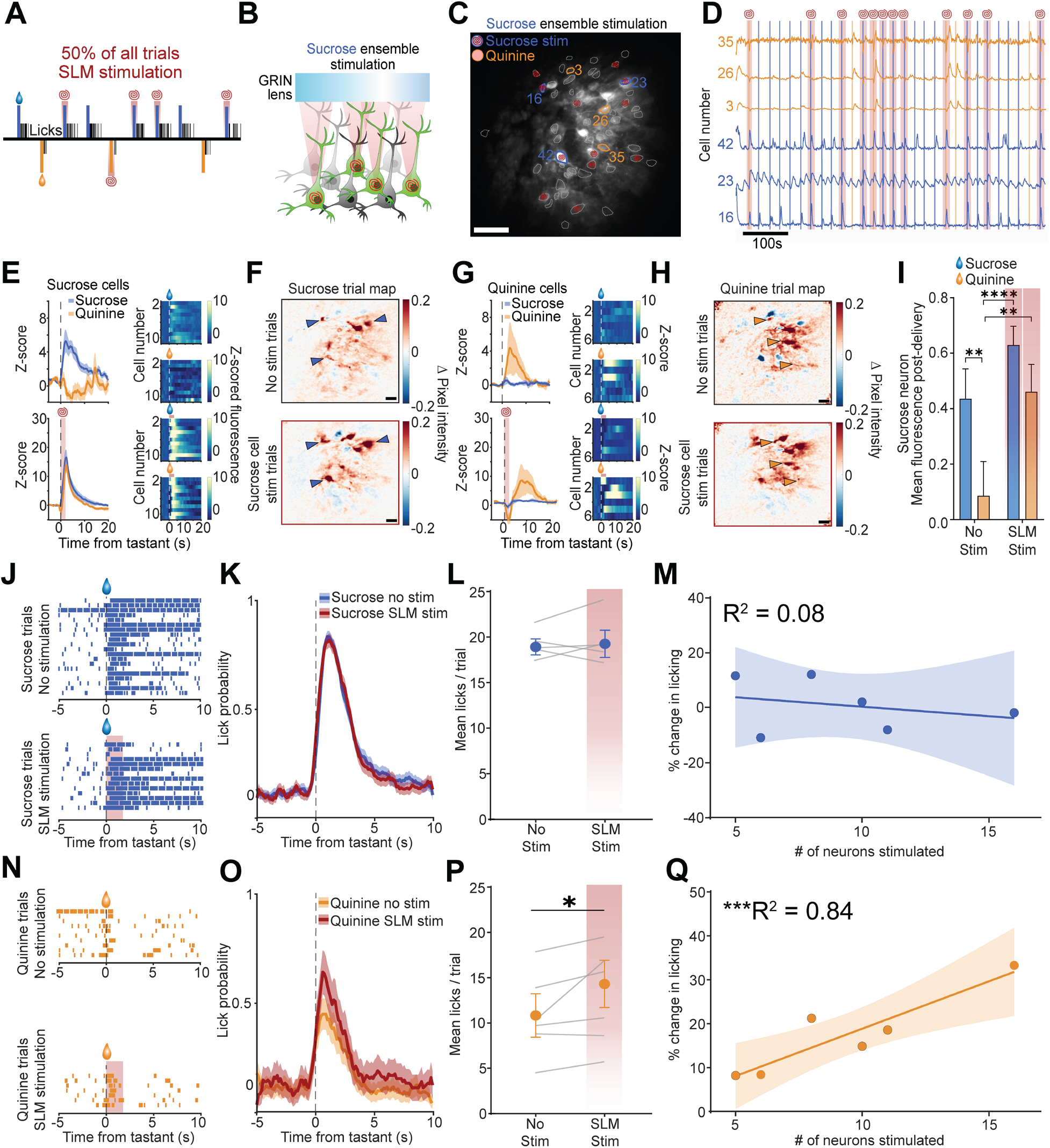
Selective stimulation of appetitive ensemble increases licking to aversive tastant. (**A**) Schematic of trial structure for SLM stimulation sessions. 70% of trials resulted in 10% sucrose (blue) delivery, while 30% of trials resulted in 2mM quinine (orange) delivery. SLM stimulation (10hz, 2s, 6-10mW power per ROI) was randomly delivered simultaneously with tastant delivery on 50% of all trials. (**B**) SLM stimulation ROIs were targeted to quinine-activated (quinine-ensemble) neurons. (**C**) Mean projection image of FOV from a single mouse. BLA neuron contours are colored according to their activity during the sucrose and quinine delivery session. Blue contours indicate neurons that were activated in response to sucrose only, while orange contours indicate neurons activated in response to quinine only. SLM spiral stimulation targets indicate the 11 sucrose activated neurons (sucrose ensemble) targeted for stimulation. (**D**) Example traces from 3 sucrose (blue) and quinine (orange) activated neurons labeled in **C**. Traces are overlayed on top of tastant delivery trials (sucrose=blue, quinine=orange) as well as trials in which SLM stimulation of the sucrose ensemble occurred (red shading). (**E**) Mean fluorescence activity of sucrose activated neurons in response to sucrose (blue) and quinine (orange) delivery on trials without stimulation (top) and trials with SLM stimulation (bottom). Dotted line indicates tastant delivery. Red shading indicates onset of quinine ensemble stimulation via SLM. (**F**) Spatial map of mean pixel intensity during the post-sucrose delivery period subtracted from mean pixel intensity immediately preceding sucrose delivery (-5s to 0s) during trials without sucrose ensemble stimulation (top) and SLM stimulation of the sucrose ensemble (bottom). Blue triangles indicate location of sucrose activated neurons identified in **C** and **D**. Red shading indicates a greater difference in activity during the post-sucrose delivery period relative to baseline. Blue shading indicates reduced activity. (**G**) Mean fluorescence activity of quinine activated neurons in response to sucrose (blue) and quinine (orange) delivery on trials without stimulation (top) and trials with SLM stimulation (bottom). Dotted line indicates tastant delivery. Red shading indicates onset of sucrose ensemble stimulation via SLM. (**H**) Spatial map of mean pixel intensity during the post-quinine delivery period subtracted from mean pixel intensity immediately preceding quinine delivery (-5s to 0s) during trials without sucrose ensemble stimulation (top) and SLM stimulation of the sucrose ensemble (bottom). Orange triangles indicate location of quinine activated neurons identified in **C** and **D**. Red shading indicates a greater difference in activity during the post-quinine delivery period relative to baseline. Blue shading indicates reduced activity. (**I**) Mean fluorescence in the post-delivery period for sucrose neurons in response to sucrose and quinine delivery on no stimulation versus SLM stimulation (two-way repeated measures ANOVA main effect of stimulation *p*<0.01, main effect of tastant *p*<0.05, stimulation X tastant interaction *p*<0.01. Sidak’s multiple comparison correction displayed). (**J**) Lick rasters from a single mouse organized by trial type and aligned to sucrose delivery on trials without sucrose ensemble SLM stimulation (top) and on trials with quinine ensemble stimulation (bottom; red shading). (**K**) Licking probability across all mice on sucrose delivery trials without stimulation (blue) and on trials with simultaneous sucrose ensemble SLM stimulation (red) (one way ANOVA, *p*>0.05). (**L**) The mean number of licks per trial per mouse on sucrose delivery trials without sucrose ensemble stimulation and on trials with sucrose ensemble SLM stimulation (red shading) (*t*(5)=0.51, *p*=0.64). (**M**) Correlation between the number of sucrose ensemble neurons stimulated via SLM and the percent change in licking on sucrose trials (No stimulation vs sucrose ensemble stimulation trials) (*R*^2^=0.08, linear regression *F*(1,4)=0.35 *p*=0.59). (**N**) Lick rasters from a single mouse organized by trial type and aligned to quinine delivery on trials without sucrose ensemble SLM stimulation (top) and on trials with sucrose ensemble stimulation (bottom; red shading). (**O**) Licking probability across all mice on quinine delivery trials without stimulation (blue) and on trials with simultaneous sucrose ensemble SLM stimulation (red). (one way ANOVA, *p*<0.05) (**P**) The mean number of licks per trial per mouse on quinine delivery trials without sucrose ensemble stimulation and on trials with sucrose ensemble SLM stimulation (red shading) (*t*(5)=3.01, *p*=0.04). (**Q**) Correlation between the number of sucrose ensemble neurons stimulated via SLM and the percent change in licking on quinine trials (No stimulation vs quinine ensemble stimulation trials) (*R*^2^=0.84, linear regression *F*(1,4)=20.45 *p*=0.01). \**p*<0.05.

We next determined whether playing back this aversive ensemble activity during trials in which an appetitive tastant (sucrose) was delivered alters consummatory behavior. Comparing sucrose consumption during trials without quinine ensemble stimulation with trials in which the aversive quinine ensemble activation was played back via SLM stimulation (**Figure 3J**) we found that lick probability was significantly reduced during stimulation trials (**Figure 4K**). Furthermore, the mean number of licks to the appetitive sucrose was significantly reduced during trials in which the quinine ensemble was activated compared to trials without SLM stimulation (**Figure 3L**). No change in licking behavior was observed when SLM stimulation was targeted to quinine ensembles in mice without ChRmine expression (**Figure S6E-G**), further supporting the conclusion the observed reduction in licking is not an artifact of SLM stimulation itself. A significant positive correlation was found between the number of quinine neurons stimulated per mouse and the reduction in licking, indicating that activating a larger aversive ensemble produces more profound reductions in consumption of sucrose (**Figure 3M**). By comparison, when quinine neurons were stimulated during quinine delivery trials within the same session, no change in the probability of licking to quinine delivery (**Figure 3N-O**) or on the mean number of licks per trial (**Figure 3P**) was detected compared to trials without stimulation. No correlation between the percentage change in licking during quinine trials and the number of neurons stimulated was detected (**Figure 3Q**). These results suggest that the endogenous activation of these neurons likely reached a ceiling and the additional activation generated by targeted photostimulation (**Figure 3I**) was not sufficient to drive any further reduction in licking. These data suggest that the neural representation of an innately aversive stimulus is hard-wired in a manner sufficient to produce aversive behavioral responding.

### Targeted photostimulation of appetitive ensemble increases consummatory behavior of aversive stimulus

We next investigated if activation of the appetitive, sucrose-responsive ensemble could produce the opposite behavioral effect (**Figure 4A-B**). Similar to quinine-ensemble stimulation, activation of the sucrose ensemble (10hz, 2s, 5-10mW power per ROI) was highly specific to sucrose and not quinine activated neurons which we identified during recordings from baseline sucrose and quinine delivery sessions (**Figure 4C-D**). Neurons that were identified as sucrose activated did not display significant activation on quinine delivery trials without sucrose-ensemble stimulation (**Figure 4E; top**). However, on trials in which sucrose ensemble stimulation (*n*=10 neurons from representative mouse) was paired with sucrose or quinine delivery, activity was not different between trial types (**Figure 4E; bottom**). Importantly, activity of quinine-responsive neurons was not impacted by photostimulation of the sucrose ensemble, as increases in activity on stimulation and non-stimulation trials were only observed in response to quinine and not sucrose delivery (**Figure 4G-H**). Across all sucrose neurons (*n*=56 total, mean per mouse=9.3, range 5-16 neurons) recorded from all mice and targeted for photostimulation (**Figure 4I**) we found that photostimulation produced a significant increase in fluorescence when paired with quinine delivery, while no significant increase in fluorescence was detected on sucrose delivery trials. No difference in mean fluorescence was detected when photostimulation was paired with sucrose or quinine. Thus, like activation of the quinine ensemble (**Figure 3**), SLM stimulation of the sucrose ensemble was specific to sucrose activated neurons, indicating that the endogenous pattern of activity evoked by sucrose delivery was able to be selectively replayed on trials in which there otherwise would have been no activity change (quinine-delivery trials). These findings strengthen the conclusion that it is possible recapitulate the spatial and temporal specificity of the endogenous response elicited by consumption of a highly appetitive tastant.

We next examined the effect of sucrose-ensemble SLM stimulation on consummatory licking behavior in response to either sucrose or quinine. In a similar manner to our observation when activating the quinine ensemble during quinine delivery trials (**Figure 3G**), photostimulation of the sucrose ensemble on sucrose delivery trials did not change licking probability (**Figure 4J-K**) or the mean number of licks per trials (**Figure 4L**) compared to trials without SLM stimulation. No significant correlation was observed between the number of neurons activated and the mean change in licking between sucrose stimulation and non-stimulation trials (**Figure 4M**). By contrast, when the appetitive sucrose ensemble was activated during aversive quinine presentation, licking was significantly increased across trials compared to trials without sucrose ensemble SLM stimulation (**Figure 4N**). Lick probability was non-significantly increased on trials in which the quinine delivery was paired with SLM activation of the sucrose ensemble relative to no stimulation trials (**Figure 4O**). However, comparing the mean number of licks per mouse in response to quinine delivery on non-stimulation trials versus trials in which the sucrose ensemble was activated, mice had a significant increase in number of licks during stimulation trials (**Figure 4P**). Additionally, a significant positive correlation was observed between the number of activated sucrose neurons and the percentage change in mean licks, supporting the concept that stimulation of greater numbers of sucrose ensemble neurons produced larger increases in quinine licking compared to trials without stimulation (**Figure 4Q**). Together these data suggest that selective recruitment of an appetitive neural ensemble is capable of biasing consummatory behavior in response to an aversive tastant toward a more appetitive-like response. In summary, this all-optical deep brain approach demonstrates that separable populations of neurons within the BLA stably encode appetitive or aversive behavioral responses, and that selective photoactivation of a small neuronal population is capable of bidirectionally influencing consummatory behavior.

## Discussion

Here we find intermixed groups of neurons within the BLA that respond reliably and selectively to appetitive stimuli, aversive stimuli, or both. At the population level, this diversity of individual neural responses allows for accurate classification of whether an appetitive or aversive stimulus was delivered, and these responses were highly stable across days, indicating stable encoding of valence within the BLA (**Figure 1**). We next established the spatial specificity of targeted photostimulation via SLM of individual BLA neurons deep in the brain through a GRIN lens for the first time, demonstrating high spatial and temporal control of targeted neurons that was robust and repeatable (**Figure 2**). Finally, we targeted BLA ensembles that responded selectively to either appetitive or aversive stimuli for photostimulation, finding that when the aversive ensemble was stimulated, licking for an appetitive tastant was reduced (**Figure 3**). By contrast, when the appetitive ensemble received photostimulation, licking for the aversive tastant was enhanced (**Figure 4**). These findings demonstrate that the neural representation of innate stimulus valence, reflected in distinct ensembles of BLA neurons with antagonistic activity profiles, is causally related to the expression of valence specific behavior.

Our findings bridge a long standing gap in our understanding of how individual BLA neuron activity directly contributes to valence processing (Tye, 2018). Numerous correlational studies have identified populations of neurons of varying sizes within the BLA that respond to appetitive and aversive stimuli as well as cues that predict them (Beyeler et al., 2018; Kim et al., 2016; Kyriazi et al., 2018; Zhang and Li, 2018). Here we identify a substantial population of neurons that respond strongly and consistently to both appetitive and aversive stimuli, suggesting that the activity of these neurons may be important for the salience of a stimulus rather than valence (Belova et al., 2007; Shabel and Janak, 2009). Due to this substantial overlap in the activity profiles of these neurons, but also their intermixed anatomical distribution, establishing whether neurons with selective activation in response to either an appetitive or an aversive stimulus simply represent the stimulus type or if they causally contribute to valence specific behavior (approach/consumption or avoidance) has not previously been possible. Here we show that these groups of BLA neurons not only have inherent and stable encoding of valence, but that their individual activation is sufficient to bias valence specific behavior. This conclusively demonstrates a causal role in controlling the expression of valence-specific behaviors by unique BLA ensembles, providing support for prior observations that BLA neurons may encode both appetitive and conditioned behavior (Kyriazi et al., 2018). In contrast to previous work which identified the BLA as instantiating positive valence while the CeA instantiated negative valence (Wang et al., 2018), we report that single neurons within the BLA contribute to both. A potential explanation for this difference is that our BLA targeting was on average more posterior (-1.5mm to -1.95mm from bregma), where it appears more overlap exists between BLA neurons that project to either sweet or bitter gustatory cortex (Wang et al., 2018). Future experiments will evaluate how conditioned stimuli are represented by these defined groups of BLA neurons and whether targeted photostimulation can alter the representation of these previously neutral stimuli.

The linear relationship detected between the number of BLA neurons photostimulated and the change in licking behavior (**Figure 3** & **4**) provides new insight into how the BLA as a whole encodes valence. This suggests that the selection of valence-specific behaviors is graded and dependent on the recruitment of stable ensembles of BLA neurons that prefer one stimulus type, as well as potentially antagonistic ensembles with the opposite response. Future experiments will evaluate whether differing concentrations of appetitive and aversive stimuli shift ensemble size, and whether the number of photostimulated neurons necessary to produce a behavioral shift depends on the perceived delta of the appetitive and aversive stimulus. If the proposed graded model of encoding valence-specific behavior exists, we expect that the more similar the appetitive and aversive stimulus is the smaller the ensemble size will become.

Prior reports have been mixed with regards to whether there is a relationship between anatomical location and whether a neuron responds selectively to appetitive or aversive stimulus (Beyeler et al., 2018; Kim et al., 2016). Here we observed no significant relationship between these variables (**Figure S3H,I**), suggesting that valence encoding neurons are found throughout the BLA.. By recording dozens of BLA neurons simultaneously we also uncover the first physiological evidence of direct antagonism of valence specific BLA neurons, which has previously been demonstrated by optogenetic activation of genetically defined BLA neurons (Kim et al., 2016). We find that a substantial proportion of neurons that respond to an appetitive stimulus are inhibited by an aversive stimulus, and vice versa (**Figure S3E-G**). This likely reflects the role of intermediate interneurons and extended microcircuits as previous reports indicate that activating discrete BLA projections produces complex network level effects including inhibition (Beyeler et al., 2018).

Here we greatly extend the depth through which targeted photostimulation of individual neurons has been achieved, which has been previously used exclusively in superficial brain structures (Adesnik and Abdeladim, 2021; Carrillo-Reid et al., 2019; Daie et al., 2021; Gill et al., 2020; Jennings et al., 2019; Marshel et al., 2019; Robinson et al., 2020). By accounting for optical aberration generated by the GRIN lens (**Figure S4**), we find that specificity of photostimulation via the SLM is qualitatively similar to what has been shown in previous experiments using cortical windows (Carrillo-Reid et al., 2019; Daie et al., 2021). Although the numbers of accessible neurons are lower than what can be achieved via a cortical window, consistent behavioral effects were observed through targeted photostimulation of ensembles responsive to either appetitive or aversive stimuli across mice (**Figure 3** **& 4**). Given the significant positive correlation we observed between the number of neurons activated and the percent change in licking behavior, we expect that more profound changes in behavior will be detected by maximizing the numbers of neurons targeted in future experiments. This could be achieved numerous ways, including using a larger GRIN lens, or by recording and stimulating multiple planes within a single mouse. Here we establish that targeted photostimulation in the deep brain is possible, and use this method to directly demonstrate a causal role for opposing BLA ensembles on the expression valence-specific behavior. In doing so, we identify a model of BLA function in the control of innate valence that can be applied to other valence domains.

## Acknowledgements

We thank Dr. Azra Suko for lab management and organization and Drs. David Ottenheimer and Adam Gordon-Fennell as well as Madelyn Hjort (all members of the Stuber lab) for assistance and input related toh head-fixed behavior. We also thank the entire Bruchas and Stuber laboratories as well as the Center for Neuroscience of Addiction, Pain, and Emotion at the University of Washington for resources, alongside critical feedback.

## Author contributions

Conceptualization: SCP, MRB, GDS

Methodology: SCP, MRB, CZZ, GDS, CP, CEP

Investigation: SCP, CZZ, CP

Visualization: SCP, CZZ, CEP, TKN, ST

Funding acquisition: MRB, GDS

Supervision: MRB, GDS

Writing – original draft: SCP, MRB

Writing – review and editing: SCP, MRB, GDS, CZZ, CP

## Data and materials availability

All data, code, and materials are available upon reasonable request.

## Figure Legends

**Figure S1.**
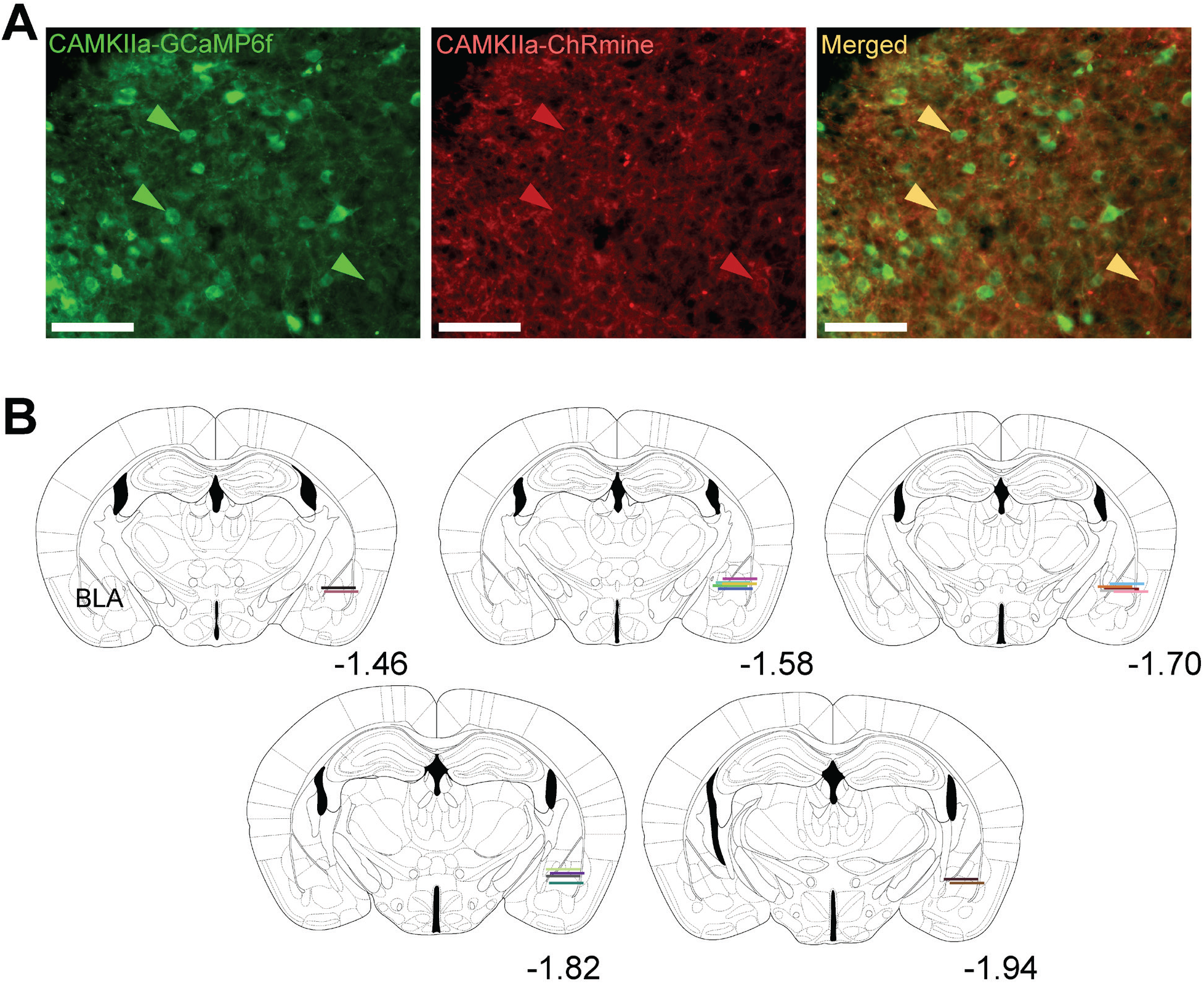
Histological confirmation of GRIN lens placements in BLA. (**A**) Histological image (20x magnification) of colocalization (right) of GCaMP6f (left) and ChRmine (middle) expression within the BLA. Arrows point to select neurons expressing both GCaMP6f and ChRmine. Scale bar = 100µm. (**B**) GRIN lens locations throughout the BLA (distance from bregma in the anterior/posterior axis in millimeters) of individual mice. Horizontal line (600µm long) indicates the bottom of the GRIN lens.

**Figure S2.**
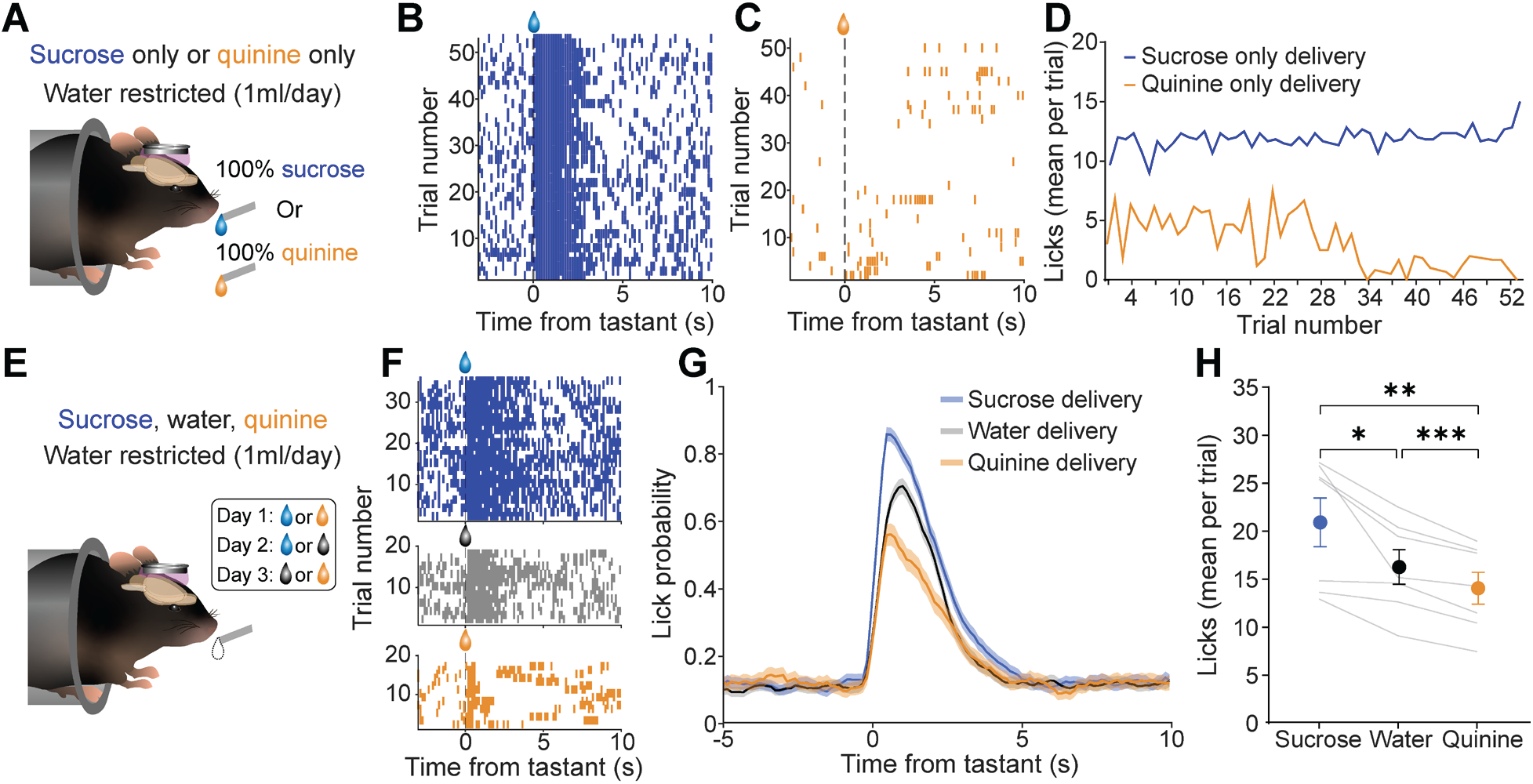
Establishing consistent appetitive and aversive behavioral response to sucrose and quinine.ss. (**A**) Schematized representation of experiment in which mice received either only 10% sucrose or only 2mM quinine on 100% of trials on separate days. (**B**) Lick rasters aligned to delivery of sucrose for a representative mouse across all trials. (**C**) Lick rasters aligned to quinine delivery for the same mouse on the quinine delivery day. (**D**) Meassn number of licks per trial across mice (*n*=6) on the sucrose only delivery day (blue) and quinine only delivery day (orange). (**E**) Schematized representation of experimental design for evaluating the difference in licking between 10% sucrose (blue), drinking water (grey), and 2mM quinine (orange). Three days each containing two tastant combinations were conducted. (**F**) Lick rasters from a single representative mouse from single sessions aligned to sucrose delivery (top; blue), water delivery (middle; grey), or quinine (bottom; orange). (**G**) Mean probability of licking across sessions and across mice on sucrose, water, or quinine trials. (**H**) Mean number of licks per trial during the post-tastant period (0 to 5s after delivery) for sucrose, water, and quinine delivery trials (one-way ANOVA; *F*(2,6)=20.36, *p*=0.003, Sidak’s multiple comparison correction sucrose vs. water (*t*(6)=3.3, *p*=0.017) sucrose vs. quinine (*t*(6)=5.6, *p*=0.001), water vs. quinine (*t*(6)=6.5, *p*=0.0006). \*\*\**p*<0.001, \*\**p*<0.01, \**p*<0.05.

**Figure S3.**
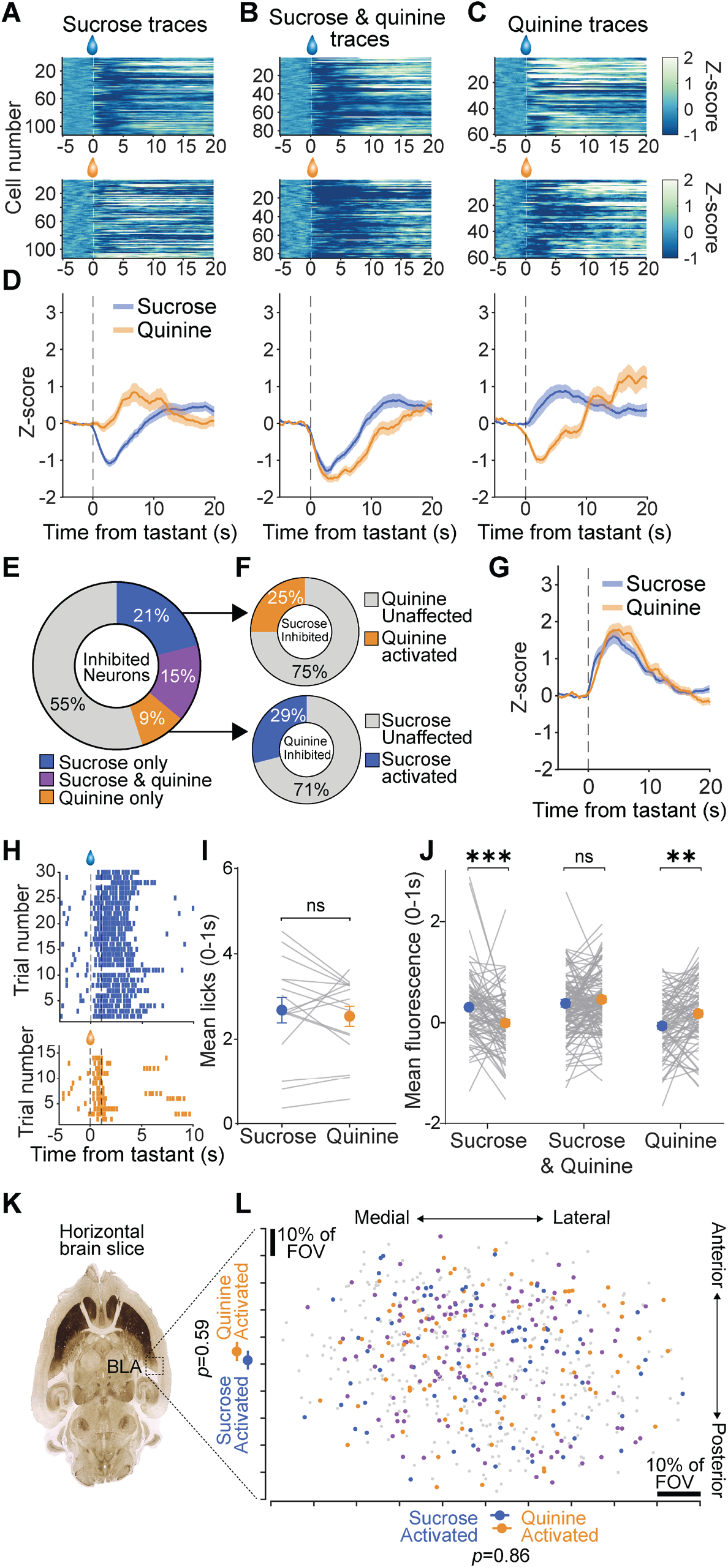
Additional characterization of tastant-evoked changes in BLA neuron activity. (**A**) Heatmaps of trial-averaged fluorescence responses of BLA neurons identified as sucrose inhibited aligned to 10% sucrose (top) or 2mM quinine (bottom) delivery. (**B**) Heatmaps of trial-averaged fluorescence responses of BLA neurons identified as sucrose and quinine inhibited aligned to sucrose (top) or quinine (bottom) delivery. (**C**) Heatmaps of trial-averaged fluorescence responses of BLA neurons identified as quinine inhibited aligned to sucrose (top) or quinine (bottom) delivery. (**D**) Mean fluorescence responses of neurons identified as sucrose inhibited (left), sucrose and quinine inhibited (middle), and quinine inhibited (right) on sucrose (blue) or quinine (orange) delivery trials. (**E**) Proportion of neurons identified as significantly inhibited in response to sucrose (blue), sucrose and quinine (purple), or quinine (orange). (**F**) Proportions of sucrose inhibited neurons that are also activated in response to quinine delivery (top; orange) and the proportion of quinine inhibited neurons that are also activated in response to sucrose delivery (bottom; orange). (**G**) Mean fluorescence responses of sucrose activated (blue) and quinine activated (orange) BLA neurons in response to sucrose and quinine, respectively (two-way repeated measures ANOVA, *p*>0.05 at all time bins). (**H**) Lick rasters from representative mouse reproduced from Figure 1E with an additional dotted line indicating 1s following delivery. (**I**) Mean number of licks during the 1s following tastant delivery. (**J**) Mean fluorescence during the 1s following tastant delivery of individual neurons classified as sucrose-only, sucrose and quinine, or quinine-only responsive. (Main effect of activated cell type [*F*(2,285)=10.10,*p*<0.0001], interaction between activated cell type and tastant delivery [*F*(2,285)=12.77,*p*<0.0001]. *Post-hoc* comparison of tastant delivery on sucrose activated neurons (*t*(285)=3.87,*p*<0.0001) and quinine activated neurons (*t*(285)=3.06,*p*=0.007), no effect of tastant delivery type on sucrose and quinine activated neuron activity *p*>0.05). (**K**) Horizontal brain slice (adapted from (Franklin and Paxinos, 2013)) showing location of BLA. (**L**) Top-down view of all BLA neurons identified, colored by selectivity (blue=sucrose activated, purple=sucrose and quinine activated, orange=quinine activated). Comparison of mean location in the M/L and A/P of sucrose activated and quinine activated neurons (two-tailed *t*-test, *p*>0.05). \*\*\**p*<0.001, *\*\*p*<0.01.

**Figure S4.**
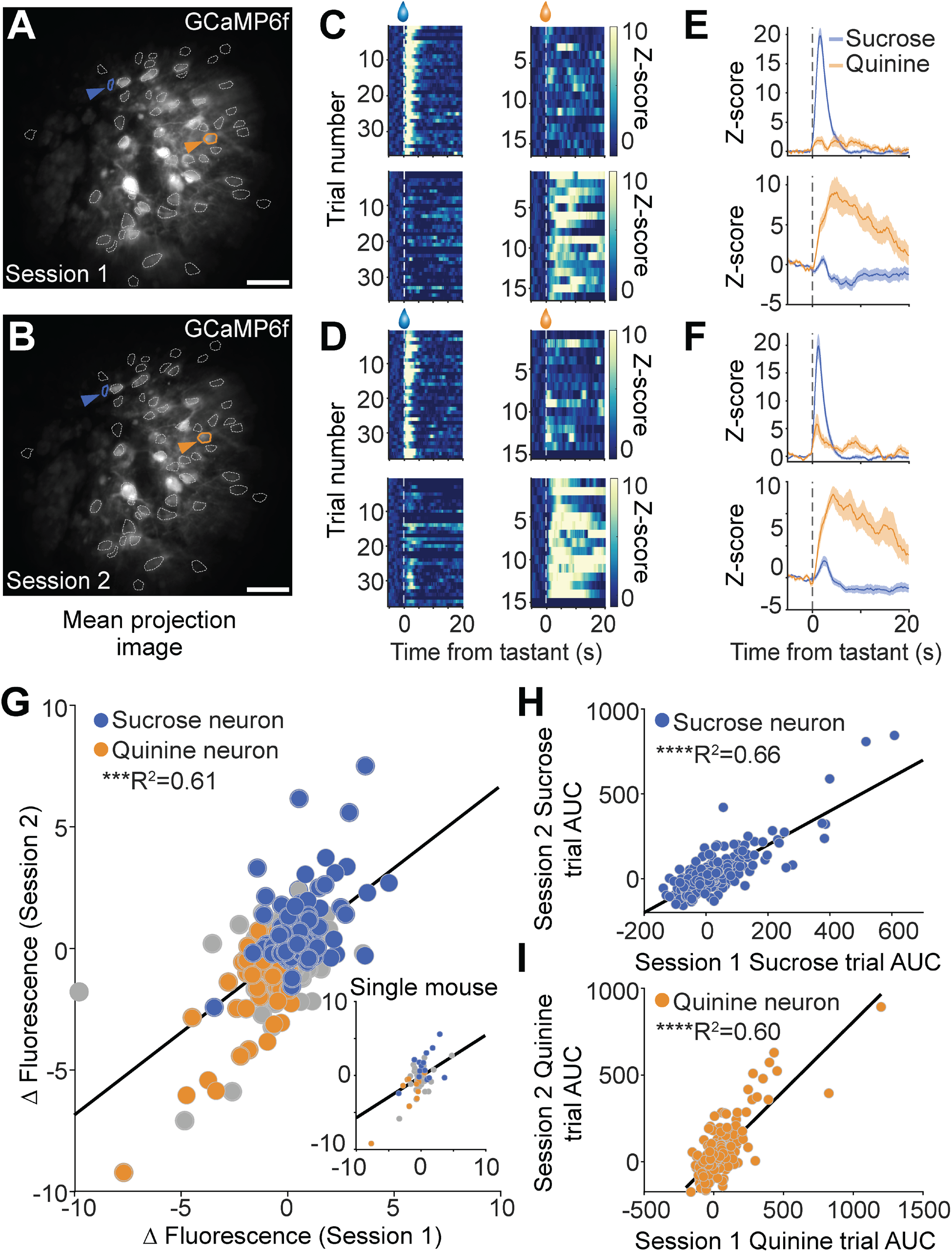
BLA neurons stably respond to appetitive or aversive stimuli. (**A**) Mean projection image of BLA neurons from one sucrose and quinine delivery session, contours identify individual putative neurons. Single blue contour and arrow indicates an example sucrose activated neuron, while the orange contour and arrow points to a quinine activated neuron. (**B**) Mean projection image of BLA neurons from a subsequent (72 hours later) sucrose and quinine delivery session. The same example neurons as in **A** are identified and colored. (**C**) Representative heatmaps of tastant-delivery aligned calcium fluorescence from individual neurons during sucrose (left) or quinine (right) delivery trials for the identified sucrose activated (top; blue) or quinine activated (bottom; orange) neuron. (**D**) Representative heatmaps of tastant-delivery aligned calcium fluorescence from the same individual neurons during subsequent sucrose (left) or quinine (right) delivery trials for the identified sucrose activated (top; blue) or quinine activated (bottom; orange) neuron. (**E**) Mean fluorescence tastant responses across trials on for the sucrose activated (top) and quinine activated (bottom) neuron on session 1 and (**F**) on session 2. (**G**) Correlation between the difference between fluorescence of individual neurons on sucrose versus quinine trials (sucrose – quinine) on Day 1 versus Day 2 (*R*^2^=0.61, *n*=303 neurons). Individual data points (neurons) are colored according to whether neurons were sucrose or quinine activated. (Inset) Fluorescence difference scores on Day 1 versus Day 2 from a single mouse. (**H**) Correlation between area under the curve (AUC) in the post-tastant delivery period for all neurons in response to sucrose delivery on Day 1 and Day 2 (*R*^2^=0.66, *n*=303). (**I**) Correlation between AUC in the post-tastant delivery period for all neurons in response to quinine delivery on Day 1 and Day 2 (*R*^2^=0.60, *n*=303). \*\*\*\**p*<0.0001, \*\*\**p*<0.001.

**Figure S5.**
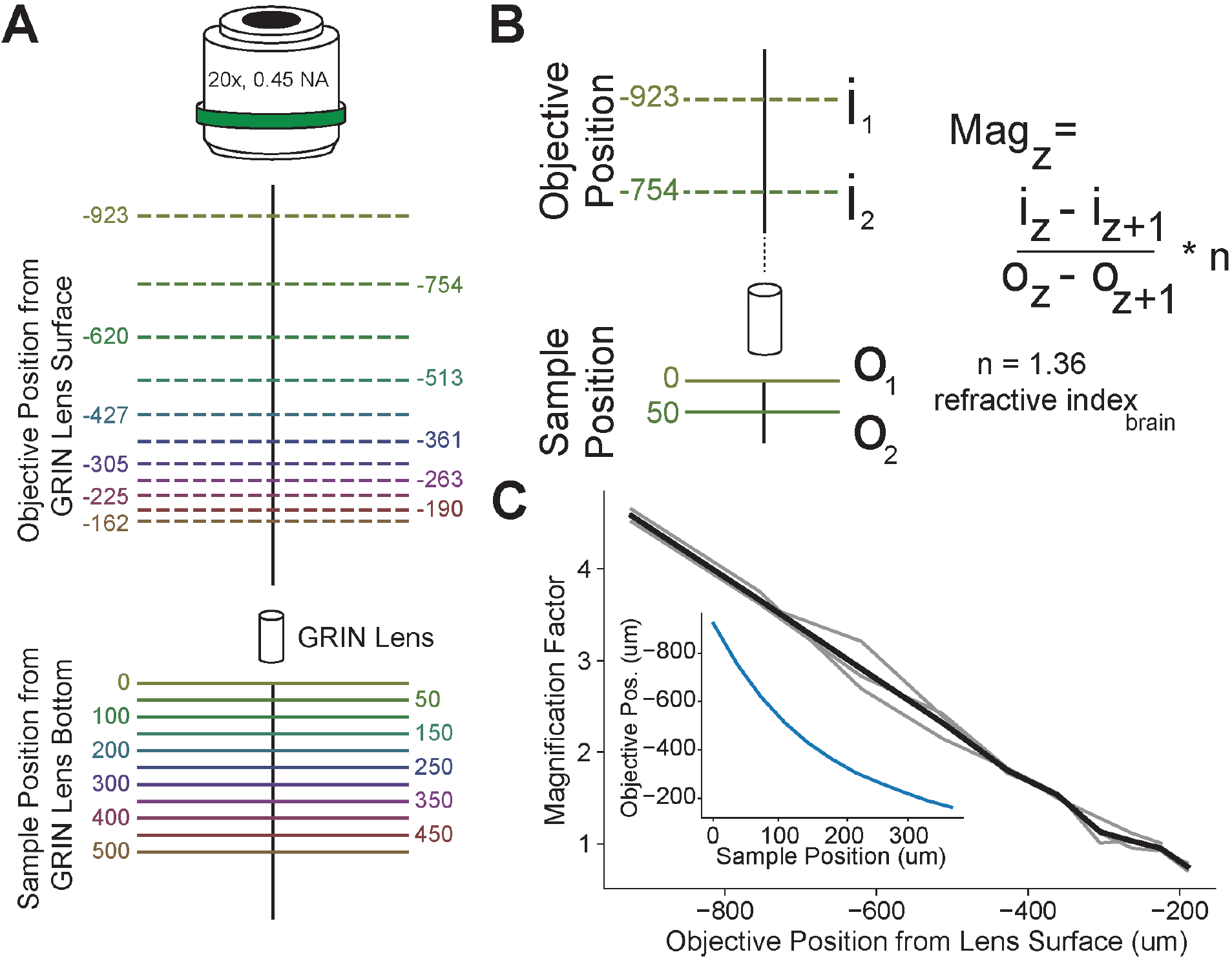
Characterization and correction of GRIN lens magnification for axial distance measurements. (**A**) Sample object positions and corresponding objective focus positions for determining degree of GRIN lens axial magnification. An object with a defining feature (chroma slide with a surface marking) was positioned at various planes relative to the bottom of a GRIN lens with a manual linear z-translation stage. The microscope objective was then refocused onto the surface of the chroma slide. Solid lines in sample space (mimicking brain tissue below the GRIN lens) correspond to the same color dotted lines in imaging space (objective focal point). Objective position measurements were averaged across data from 3 different GRIN lenses. (**B**) Example of axial magnification calculation. Using the corresponding sample and objective position data measured in **A**, we calculated a magnification factor, which consisted of dividing the distance between two objective focal planes by the distance between the corresponding sample position planes. Finally, this value was multiplied by the approximate refractive index of brain tissue (1.36). i_z_ : coordinate of plane Z in image space (ie. objective position), o_z_ : coordinate of plane Z in image space (ie. sample position). (**C**) Magnification factor as a function of objective focal position. Gray lines indicate data from individual GRIN lenses (n=3), Black line represents the average across GRIN lenses. Inset: Alternate representation of the data: objective position as a function of the corresponding sample position.

**Figure S6.**
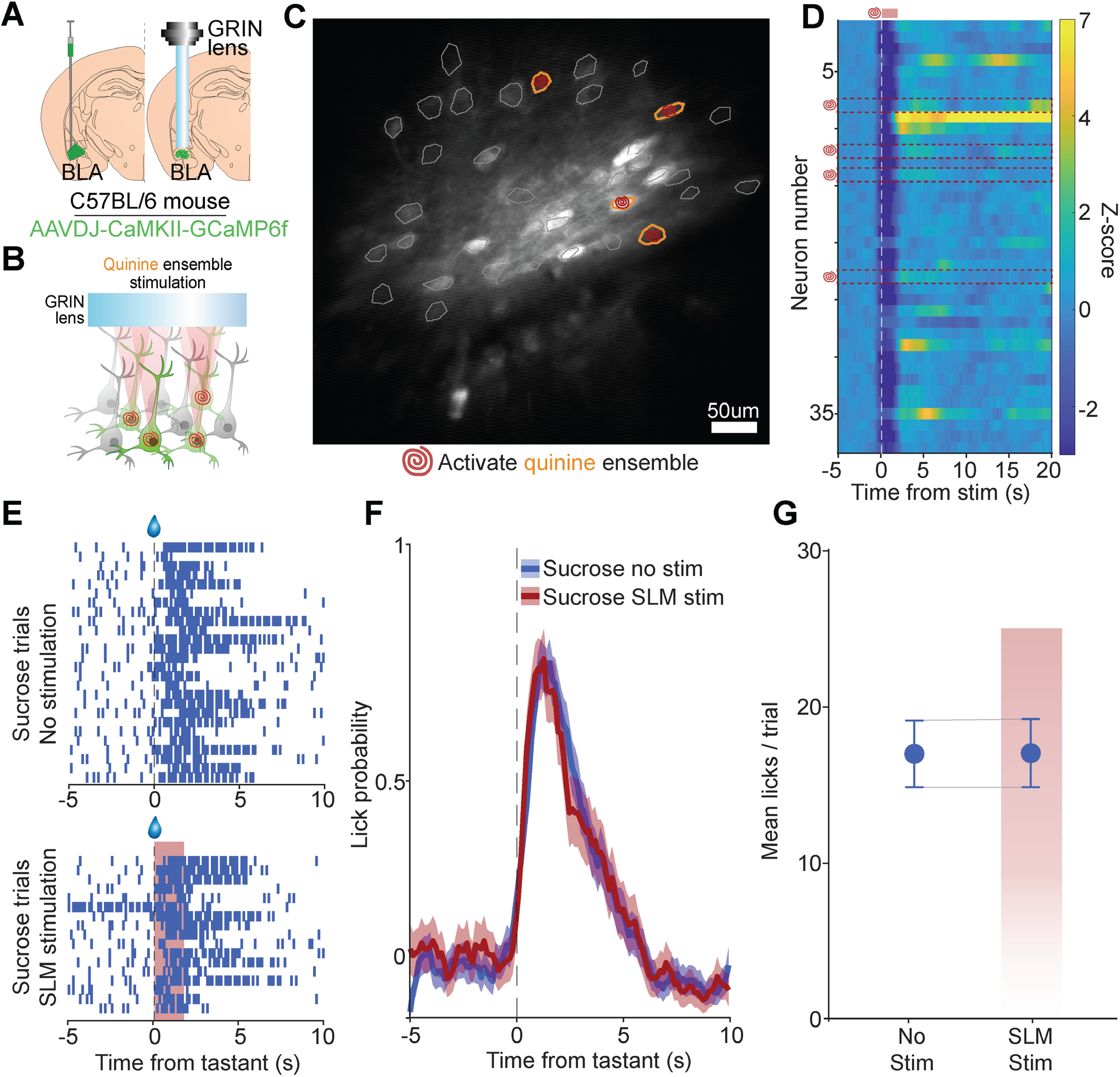
SLM stimulation of quinine-ensemble neurons in ChRmine-negative mice does not alter licking on sucrose trials. (**A**) Schematic of surgical procedure injecting an AAV into the BLA of C57BL/6 mice to express GCaMP6f under the CAMKII promotor. No viral vector expressing ChRmine was injected. (**B**) Experimental design targeting quinine ensemble neurons for SLM activation (10hz, 2s, 6-10mW power per ROI) during session in which mice receive random presentations of sucrose and quinine. (**C**) Mean projection image of FOV from a single ChRmine-mouse during the quinine ensemble stimulation session. Quinine ensemble neurons are identified by orange contours and spiral stimulation targets mark ROIs for SLM stimulation. (**D**) Heatmap of trial-averaged fluorescence across all ChRmine-neurons during quinine ensemble stimulation session aligned to SLM stimulation onset. Red dashed outlines and spirals indicate quinine neurons targeted for SLM stimulation. (**E**) Lick rasters from a representative ChRmine-mouse aligned to sucrose delivery on trials without SLM stimulation (top) and trials with SLM stimulation (bottom; red shading). (**F**) Mean licking probability of on sucrose delivery trials without quinine ensemble stimulation (blue) and with quinine ensemble SLM stimulation (red). (**G**) The mean number of licks per trial per mouse on sucrose delivery trials without quinine ensemble stimulation and on trials with quinine ensemble SLM stimulation (red shading) (*n=*2, *t*(1)=1.0, *p*=0.50).

**Extended Data Movie 1. Stimulation of quinine ensemble during task**.

*In vivo* two-photon calcium imaging of BLA neurons during a session in which quinine ensemble neurons were targeted for holographic photostimulation. Video depicts selected quinine ensemble neurons (red) targeted for photostimulation (10hz, 2s duration, 5ms pulse width) during the presentation of either sucrose or quinine. Video acquired at 7.5hz and played back at 13x speed. Initial targeted photostimulation is slowed down to 3x speed.

## STAR Methods

| REAGENT or RESOURCE | SOURCE | IDENTIFIER |
| --- | --- | --- |
| Bacterial and virus strains |  |  |
| AAV5-CAMKIIa-GCaMP6f-WPRE-SV40 | Penn Vector Core |  |
| AAV8-CAMKIIa-ChRmine-mScarlet-Kv2.1-WPRE | Stanford Vector Core | GVVC-AAV-194 |
| Chemicals, peptides, and recombinant proteins |  |  |
| VECTASHIELD Hardset Antifade Mounting Medium with DAPI | Vector Laboratories | CAT#H-1800 |
| Sucrose, 99% | Fisher Scientific | AAA155830C |
| Quinine hydrochloride dihydrate | Sigma-Aldrich | Q1125 |
| Experimental models: Organisms/strains |  |  |
| C57BL/6J | Jackson Laboratory | 000664 |
| Software and algorithms |  |  |
| FIJI/ImageJ | NIH | https://fiji.sc/ |
| Prairie View | Bruker |  |
| MATLAB | Mathworks | https://www.mathworks.com/products.html |
| Python 3.0 | Python Software Foundation | https://www.python.org/ |
| SIMA | (Kaifosh et al., 2014) | http://www.losonczylab.org/sima/1.3.2/ |
| NAPE Calcium imaging pipeline | Github | https://github.com/zhounapeuw/NAPE_imaging_analysis |
| PRISM 8 | GraphPad | https://www.graphpad.com/ |
| Illustrator CS8 | Adobe | https://www.adobe.com/products/illustrator.html |
| Other |  |  |
| ProView GRIN lens (0.6mm x 7.3mm) | Inscopix | 1050-004608 |
| Ultima 2P Plus w/ NeuraLight 3D Spatial Light Modulator | Bruker | https://www.bruker.com/en/products-and-solutions/fluorescence-microscopy/multi-photon-microscopes/ultima-2pplus.html |
| Insight X3 Tunable Ultrafast Laser | Spectra-Physics | https://www.spectra-physics.com/en/f/insight-x3-tunable-laser |
| Spirit High Power Femtosecond Lasers | Spectra-Physics | https://www.spectra-physics.com/en/f/spirit-femtosecond-laser |

### Animals

All procedures were carried out in accordance with the guidelines for the care and use of laboratory animals from the NIH and with approval from the University of Washington Institutional Animal Care and Use Committee (IACUC). Male and female C57BL/6 mice were used for all experiments. Mice in all cohorts were approximately 3-5 months old at the time of surgery. Mice were housed on a reverse light/dark cycle (lights on: 9:00PM, lights off 9:00AM).

### Stereotactic surgery

For all surgeries, mice were anesthetized using 5% isoflurane mixed with oxygen and maintained at 1-2% isoflurane maintenance for the duration of the surgery. Mice were secured on a small-animal stereotactic instrument (Kopf Instruments, Tujunga, CA USA). Body temperature was maintained throughout surgery at 37°C using an infrared heating pad (Kent Scientific, Torrington, CT USA). Ophthalmic ointment was immediately applied to both eyes once mice were anaesthetized to prevent drying. Three rounds of betadine and alternating 70% ethanol wipes were applied to the scalp prior to an incision being made down the midline suture. A single skull screw was placed just in front of the lambdoid suture at a depth that did not penetrate the skull.

After skull screw placement, a 1mm drill-bit was used to create a craniotomy above the BLA (coordinates A/P: -1.45, M/L: -3.22, D/V: -4.55). The D/V coordinate was measured relative to skull surface above the BLA, while A/P and M/L were measured from an interpolated bregma. 700nl of a 1:3 mixture of opsin (AAV8-CAMKIIa-ChRmine-mScarlet-Kv2.1-WPRE; Stanford Viral Vector Core, Titer 2.3×10^13^) and GCaMP6f (AAV5-CAMKIIa-GCaMP6f-WPRE-SV40, Penn Viral Vector Core, Titer 1.31×10^13^) was injected using a fixed-needle Hamilton Neuros syringe (Hamilton Company, Reno, NV USA) and an infusion pump (World Precision Instruments, Sarasota, FL USA) at a rate of 100 nl/min. Immediately after viral injection, a GRIN lens (Inscopix, ProView 7.3mm length, 0.6mm diameter) was lowered directly above the viral injection target at the same coordinates. The GRIN lens was lowered steadily until the D/V approached - 3.55mm from skull surface, at which point it was lowered slowly at a rate of 100 µm/min until reaching final depth of -4.55mm. The skull was then dried and a layer of all purpose krazy-glue (Krazy Glue, Columbus, OH USA) was applied to the skull. After this application, a layer of black dental acrylic (Lang Dental, Wheeling, IL USA) was added to the skull and sides of the GRIN lens and built up into a secure base. A custom machined steel head-ring was then placed on top of the acrylic base such that the GRIN lens was in the center of the ring. The ring was then secured in place with another layer of dental acrylic. Animals were injected with slow-release Carprofen (5 mg/kg) prior to surgery to act as a post-surgery analgesic. Animals were monitored daily for 7 days following surgery.

### Water restriction protocol

Mice recovered from surgery for a minimum of 4 weeks before beginning water restriction. Mice were water-restricted by giving them access to water only once per day in order to properly motivate consumption of sucrose, quinine, or water in behavioral assays. Starting 1 week prior to behavioral testing, the mice were given 1.0 mL of water in a 2×2 cm paper cup in their home cage each day at approximately 3-4pm. The mice promptly consumed the water while monitored by the researcher to assure they consumed most of the water. Animals were also weighed routinely to ensure they were at 85-90% of their body weight while water-restricted. No health issues related to dehydration occurred throughout the water-restriction period. After the behavioral testing concluded, water-restriction was discontinued and the mice were given *ad-libitum* access to water in their home cages again.

### Sucrose and quinine behavioral paradigm

After recovery from surgery, the mice were water restricted (see Water Restriction Protocol) and trained to lick 10% sucrose solution while immobilized and head-fixed. Head-fixation was accomplished by scruffing the mice, putting their back end into a 70 mL conical tube, and then placing the tube with the mouse into a custom headstage (Namboodiri et al., 2019). All mice were habituated to the head-fixation and lick training with 10% sucrose for 4 days. After habituation, a custom built Arduino controlled liquid delivery system was used to randomly deliver either 10% sucrose (70% of trials, pseudorandom) or 2mM quinine (30% of trials, pseudorandom) via distinct solenoids, tubes, and lick spouts. The licks spouts were positioned in a triangular formation such that liquid was accessed from the same point regardless of trial type. The intertrial interval was 20-25 seconds. Discrete lick events were detected and recorded using a contact lickometer circuit. TTLs reflecting each event (tastant delivery and licks) were sent directly to the data acquisition system on the two-photon microscope to ensure accurate tracking of both behavior and imaging data. The behavioral testing was conducted in the dark.

### Sucrose-only and quinine-only paradigm

After undergoing lick training and the sucrose and quinine behavioral paradigm listed above, a subset of mice underwent a sucrose-only paradigm for 1 day. The mice were head-fixed as described above. Testing was performed with a 20 minute session with 100% of trials delivering 10% sucrose. Delivery of sucrose and recording of licks was the same as described in the sucrose and quinine behavioral paradigm above.

Following the sucrose-only paradigm listed above, the same mice underwent a quinine-only paradigm for 1 day. The mice were head-fixed as described above. The test consisted of a 20 minute session with 100% of the trials delivering 2mM quinine. Delivery of quinine and recording of licks was the same as described in the sucrose and quinine behavioral paradigm above.

### Sucrose, water, and quinine behavioral paradigm

To compare licking of the appetitive tastant sucrose and aversive tastant quinine to regular drinking water, a 3 day paradigm was established. On the first day, the lick-trained mice first underwent a head-fixed sucrose and water test, consisting of a 30 minute session where they were presented with a droplet of either 10% sucrose or water. The percentage of sucrose and water trials were 70% and 30%, respectively, which were presented pseudorandomly. On the second day, the mice were offered a droplet of either 2mM quinine solution or water during the 30 minute session. The percentage of water and quinine trials were 70% and 30%, respectively, which were presented pseudorandomly. Conditions of delivery of the solutions and recordings of the licks were the same as described above.

### General two-photon imaging

4 to 6 weeks after surgery mice were habituated to head-fixation for at least 4 consecutive days. Head-fixation was achieved by securing the circular head-ring into a metal clamp attached to a custom head-stage while the body of the mouse was lightly restrained in a 70ml conical tube. During habituation, mice were placed underneath the two-photon microscope for 5 minutes and given access to random presentations of 10% sucrose to familiarize mice with the procedure. Following habituation, combined two-photon imaging and behavior sessions were conducted. GCaMP6f imaging was acquired via an Ultima 2P Plus with the Neuralight 3D Spatial Light Modulator (SLM) (Bruker Fluorescence Microscopy, Middleton WI, USA) two-photon microscope using Prairie View Software (Bruker Fluorescence Microscopy, USA). Individual frames were acquired at 30Hz using a galvo-resonant scanner and a 4 times temporal average was performed on-line for an effective frame rate of 7.5Hz with a resolution of 512px x 512px. We used a long working distance 20x air objective designed for infrared wavelengths (Olympus, LCPLN20XIR, 0.45 numerical aperture (NA), 8.3mm working distance) combined with an Insight X3 Tunable Ultrafast Laser at 920nm (Spectra-Physics, Santa Clara, CA USA). The microscope was also equipped with an electrically tunable lens (ETL) in the resonant scan path. The ETL provides dynamic positioning of the imaging focal plane and is decoupled from the photostimulation laser path, enabling simultaneous imaging and photoactivation at different Z-planes. The same imaging FOV was identified by carefully measuring the distance from the top of the GRIN lens to the field of view from day to day.

### Photostimulation during behavior

For targeted photostimulation, the same microscope and acquisition system (Bruker) was used with a second laser path consisting of a 1040nm high power femtosecond pulsed laser (Spirit One, Spectra-Physics), spatial light modulator (512×512 pixel density) to generate multi-point stimulation montages, and a pair of galvanometer mirrors for generating spiral patterns of the montage (Yang et al., 2018). Neurons were selected for targeted photostimulation based on their selectivity for either sucrose or quinine (see **Tastant related activity classification**) from a prior session. During the photostimulation session, the correct FOV was identified and a 128 frame average image was generated in order to clearly highlight all neurons. Using the selectivity map (e.g. **Figure 1F**) as a reference, spiral stimulation targets (15-20µm diameter) were manually placed on top of GCaMP6f-expressing neurons selective for a particular tastant (sucrose or quinine). Laser power was adjusted based on the number of spiral stimulation targets (6-10mW of stimulation per target) for each individual animal and session.

Using our custom Arduino code for controlling behavioral parameters, a TTL was randomly delivered to the Bruker system to elicit holographic photostimulation (20hz, 2s, 5ms pulse width) of select ensembles on 50% of tastant delivery trials (either sucrose or quinine delivery). Tastant delivery and stimulation occurred simultaneously for all experiments. With the exception of randomly delivered photostimulation, all other aspects of the behavioral paradigm remained the same. Due to the manual nature of spiral stimulation placement, experimenters were not blind while photostimulation sessions were conducted.

### Lateral and axial resolution photostimulation

To measure the physiological resolution of 2-photon holographic photostimulation, two sets of experiments were performed for lateral and axial dimensions. For estimating lateral resolution, an isolated target cell was initially identified. A stimulation protocol with the same temporal and spatial parameters as our main experiments was set up, with the exception that the SLM mask targeted three ROIs and five stimulation trials were collected. One of these ROIs was positioned over the target cell and the other two were positioned outside of the GRIN lens FOV. The reason for this motif was to properly engage the SLM, as single ROI stimulation would not utilize the SLM. We then repeated the stimulation protocol several times with the SLM mask shifted laterally by the size of a cell (20µm) for each iteration. The data were then analyzed to examine response magnitude as a function of stimulation ROI distance from the target cell.

For measuring axial resolution, an isolated target cell was again identified. The same protocol as outlined for the lateral resolution experiments were performed up until shifting the ROI mask. Instead, in order to move the stimulation ROIs in the Z-dimension, the microscope Z position was offset by a predetermined distance. This step would also offset the imaging plane; however we aimed to record target cell activity in response to out of plane stimulation. Accordingly, we compensated for this imaging plane displacement by offsetting the microscope’s ETL in the opposite direction of the microscope Z movement. We then repeated this stimulation protocol several times with the microscope Z positioned at randomized distances and corresponding ETL offsets. The data were then analyzed to examine response magnitude as a function of stimulation ROI distance from the target cell.

### Histological confirmation of virus and GRIN lens targeting

Mice were transcardially perfused with 10% formalin and phosphate buffered saline (1X PBS). Immediately after perfusion, heads (with implants intact) were placed into 10% formalin for 24h for post-fixation after which brains were removed and transferred to a 30% sucrose (in 1X PBS) solution. Brains were frozen and 35um sections were cut on a cryostat in 1:6 series. 1 series was mounted, counterstained with DAPI mounting media (Fluoroshield, Sigma-Aldrich) and imaged at 20x resolution on epifluorescence (Leica Microsystems, Wetzlar, Germany) to visualize virus and implant sites. If GRIN lens implants were determined to be outside the BLA (**Figure S1**), mice were excluded from the study.

### Two-photon imaging data processing

Individual .tif files from a single recording session were collected at 7.5Hz and combined into HDF5 format using custom code. HDF5 files were then imported into the Sequential IMaging Analysis (SIMA) Python package (Kaifosh et al., 2014) as previously described for motion correction in the x-y plane (Namboodiri et al., 2019). Following motion correction, the motion corrected HDF5 file as well as the mean projection image across all motion corrected frames were imported into Fiji is just ImageJ (Fiji, version 1.52) (Schindelin et al., 2012) for selecting neuronal regions of interest (ROIs). ROIs were manually drawn by a trained observer using the polygon selection tool. Care was taken to capture only somatic ROIs and to minimize any potential overlap between multiple soma. ROIs were then imported into SIMA for signal extraction. These aforementioned preprocessing steps were facilitated by the NAPECA analysis pipeline that wraps around the SIMA package (https://github.com/zhounapeuw/NAPE_imaging_analysis). Across sessions the same ROI file was used with minor adjustments depending on slight translation or rotation of the field of view (FOV) across sessions. Tracking across sessions was highly reliable (**Figs. 2,4,5**).

Continuous SIMA extracted fluorescence signals for each neuron were then normalized to the mean and standard deviation from the entire session (Z-score). This normalized fluorescence was used for all down-stream analyses.

### Tastant related activity classification

In order to perform unbiased classification of an individual neuron’s responsiveness (activated, inhibited, or unaffected) to a tastant delivery (e.g. sucrose or quinine delivery) we adapted a previously used strategy (Jimenez et al., 2018; Piantadosi et al., 2022). For each individual cell, raw calcium traces 3 seconds prior to grooming onset and 10 seconds after grooming onset (188 total samples at 7.5 Hz, 133ms per frame) were shuffled in time for each sample (200x) removing any temporal information that was previously in each trace but maintaining the variance within each trial. This shuffle was then performed 1000 times per cell to obtain a null distribution of tastant delivery associated calcium activity. A cell was considered responsive to a particular tastant onset if its average behavioral event Z-normalized calcium fluorescence amplitude between 0s before delivery to 3s after delivery exceeded a 1 standard deviation threshold from the null distribution.

To categories neurons as sucrose-only, quinine-only, or sucrose and quinine responsive, logical indexing was used. Thus, a sucrose-only activated neuron had significantly increased activity in response to sucrose without a significant increase to quinine. Likewise, a quinine-only activated neuron had significant increase in activity in response to quinine without a significant increase to sucrose. Sucrose and quinine activated neurons had significant increases in activity in response to both sucrose and quinine delivery. For inhibited neurons, the same indexing rule applied.

### Population decoding of tastant type

To investigate whether delivery of sucrose or quinine could be uniquely identified only by the simultaneous activity of BLA neurons, a linear support vector classifier (SVC) model (cost = 0.8) was created. The linear SVC model was trained and tested on the data for each animal individually. The class of the tastant condition and the magnitude of the reward response (0-5 s from onset of tastant delivery) of every neuron for a given trial was used as input for the model. 75% of the trials were used to train the linear SVC and 25% were used for cross-validation. The accuracy of the model during cross-validation for each animal is reported in the Results and **Figure 1**. The linear SVC model was created in Python using the *Scikit-learn* library.

### Anatomical distribution of tastant-selective neurons

To investigate the relationship between valence selectivity of BLA neurons and relative anatomical position, we calculated the relative position of tracked neurons in the focal plane during 2-photon calcium imaging sessions. The 2-photon resonant scanner generated raw frames that were 512×512 pixels. The mean x/y position of the manual drawn ROI was generated for each neuron in MATLAB. Medial-lateral and anterior-posterior axes were inferred based on the orientation of the animal being orthogonal to the objective lens during the imaging and behavior session. Due to magnification produced by the GRIN lens in the axial dimension and imprecision in determining the precise histological location of the imaged plane of neurons *ex vivo*, this dimension was not taken into account in our anatomical analysis.

### Statistical analysis of fluorescence data

Analysis of continuous fluorescence traces comparing activity of neurons in response to sucrose or quinine were carried out using two-way repeated measures ANOVA. When a significant tastant X time interaction was detected, post-hoc comparisons of individual time bins were conducted using Sidak’s multiple comparison correction. Unless otherwise noted in the main text, mean post-delivery fluorescence was calculated for each individual neuron (trial-averaged fluorescence) using a window from 0 to 5s post-tastant delivery. To compare mean fluorescence of individual neurons, again a two-way repeated measures ANOVA was used with Sidak’s multiple comparison correction if a significant interaction was observed.

To probe for stability of valence encoding, Pearson’s correlations were used. For assessing the stability of fluorescence amplitude, a difference score (Sucrose – quinine) was conducted using the trial-averaged fluorescence responses for each individual neuron. Thus, for each session a single value (Δ Fluorescence) was computed for each neuron for each session. A positive value indicates a sucrose-selective neuron, while a negative value indicates a quinine-selective neuron. These difference scores were then correlated between the two recording sessions. To assess the consistency of the temporal dynamics of the calcium signal across sessions in response to sucrose or quinine, the area under the curve (AUC) of the trial-averaged fluorescence was obtained for each individual neuron using the *trapz()* function in MATLAB. These AUC values were then correlated across sessions. GraphPad Prism 8.0 (GraphPad, La Jolla, CA USA) was used for all statistics and correlations. Unless otherwise noted, alpha was set to 0.05 and plotted data represent mean ±standard error of measurement (SEM).

### Statistical analysis of licking behavior

Mean number of licks on sucrose or quinine trials (and on photostimulation trials versus no stimulation trials) was analyzed using a paired two-tailed independent samples t-tests. For comparing the probability of licking in response to sucrose or quinine delivery two-way repeated measures analysis of variance (ANOVA) was used. Main effects and interactions are reported and in the case of significant interactions, post-hoc comparisons of significant time bins were made using Sidak’s multiple comparison correction. Two-way repeated measures ANOVAs were also used for analyzing the effect of targeted photostimulation on licking probability in response to either sucrose or quinine. GraphPad Prism 8.0 (GraphPad, La Jolla, CA USA) was used for all statistics and correlations. Unless otherwise noted, alpha was set to 0.05 and plotted data represent mean ±standard error of measurement (SEM).

